# Recovery of the gut microbiota after antibiotics depends on host diet and environmental reservoirs

**DOI:** 10.1101/717686

**Authors:** Katharine M. Ng, Andrés Aranda-Diaz, Carolina Tropini, Matthew Ryan Frankel, William W. Van Treuren, Colleen O’Laughlin, Bryan D. Merrill, Feiqiao Brian Yu, Kali M. Pruss, Rita Almeida Oliveira, Steven K. Higginbottom, Norma F. Neff, Michael A. Fischbach, Karina B. Xavier, Justin L. Sonnenburg, Kerwyn Casey Huang

## Abstract

That antibiotics alter microbiota composition and increase infection susceptibility is well known, but their generalizable effects on the gut commensal community and dependence on environmental variables remain open questions. Here, we systematically compared antibiotic responses in gnotobiotic and conventional mice across antibiotics, microbiotas, diets, and housing status. We identify remarkable resilience, whereby a humanized microbiota recovers before drug administration ends, with transient dominance of resistant *Bacteroides* and taxa-asymmetric reduction in diversity. In other cases, *in vitro* sensitivities were not predictive of *in vivo* responses, underscoring the significance of host and community contexts. A fiber-deficient diet exacerbated collapse of the microbiota and delayed recovery, despite the presence of a similar core community across diets at the point of maximal disturbance. Resilience to a second ciprofloxacin treatment was observed via response reprogramming, in which species replacement after ciprofloxacin treatment established resilience to a second treatment, and also through cross housing transmission. Single-housing drastically disrupted recovery, highlighting the importance of environmental microbial reservoirs and suggesting sanitation may exacerbate the duration of antibiotic-mediated disruption. Our findings highlight the ability of the commensal microbiota to deterministically adapt to large perturbations, and the translational potential for modulating diet, sanitation, and microbiota composition during antibiotics.

## Introduction

The gut microbiota performs functions crucial to the host, guiding development of the immune system, tuning inflammation, and establishing resistance against enteric pathogens (Ivanov et al., 2008; Mazmanian et al., 2008; Stecher et al., 2007). Antibiotic treatment perturbs community structure and function, often resulting in prolonged pathogen susceptibility (Doorduyn et al., 2006; Pavia et al., 1990; Stecher et al., 2007). Moreover, antibiotic treatment has long-lasting effects on physiological processes such as adiposity, insulin resistance, and cognitive function (Cho et al., 2012; Cox et al., 2014; Frohlich et al., 2016; Hwang et al., 2015). The ability to accurately predict how a given antibiotic will affect a community may reveal control mechanisms for these wide-ranging effects.

The manner and extent to which complex communities respond to and recover from antibiotics is not consistent. For example, while treatment with the aminoglycoside streptomycin shifted the balance between the two dominant phyla in the mouse gut strongly toward a Bacteroidetes-dominated microbiota in one study (Thompson et al., 2015), Firmicutes dominated in others (Ng et al., 2013; Stecher et al., 2007). The fluoroquinolone ciprofloxacin elicited highly individualized responses in humans (Dethlefsen et al., 2008; Dethlefsen and Relman, 2011). Many factors across studies and between individual hosts could account for variable outcomes, motivating a more systematic analysis of treatment outcomes. Moreover, few studies have focused on the dynamics of recovery after antibiotic treatment despite the relevance to pathogen susceptibility (Doorduyn et al., 2006).

Antibiotic response may depend on a wide range of environmental perturbations that multifactorially impact the microbiota and the intestinal environment. Removal of microbiota-accessible carbohydrates (MACs) from the mouse diet thins the mucosal layer surrounding the gut epithelium (Desai et al., 2016; Earle et al., 2015), increasing sensitivity to certain pathogens (Desai et al., 2016), and reduces microbiota transmission between generations (Sonnenburg et al., 2016). This dietary shift also predisposes mice to infection with *Clostridium difficile* (Hryckowian et al., 2018). Fecal microbiota transplants can replenish reservoirs of commensals in dysbiotic communities to resolve recurrent infections of *C. difficile* in humans (van Nood et al., 2013), and are also effective at restoring microbiota species after diet- (Sonnenburg et al., 2016) or osmotic diarrhea-mediated loss (Tropini et al., 2018), suggesting the importance of general environmental microbial reservoirs for commensal recovery. In some humans, a second treatment of ciprofloxacin resulted in larger disruption of the gut community than a treatment 6 months earlier (Dethlefsen and Relman, 2011). The factors that lead to community disruption after a second treatment, and the microbiota response to combinations of antibiotic treatment with other gut perturbations, are underexplored and likely critical for understanding how antibiotics reshape the human gut.

Can the wealth of *in vitro* data on the antibiotic susceptibility of model organisms be used to predict the response of communities, particularly within a host? A recent screen of 40 commensals abundant in humans across a range of antibiotics and human drugs demonstrated that sensitivity profiles can be highly species-specific (Maier et al., 2018), indicating that phylogeny is not generally useful for inferring the response of related species. Moreover, growth rate can greatly affect minimum inhibitory concentrations (MICs) *in vitro* (Evans et al., 1991), suggesting that the spatially heterogeneous and dynamic nutrient environment of the gut further confounds predictions even in simple defined communities. In a few cases, mechanisms have been identified by which one species can alter the sensitivity of another species, for instance through drug metabolism (Onderdonk et al., 1979). Last, community interactions may perturb sensitivity in a widespread manner. For example, the cell wall-targeting antibiotic vancomycin, which primarily affects only Gram-positive species *in vitro* due to the permeability barrier of the Gram-negative outer membrane, nevertheless reduces the levels of Gram-negative species in the mouse gut (Ivanov et al., 2008). These factors motivate detailed analyses of the dynamics of species-level abundances throughout antibiotic treatment and comparisons with *in vitro* sensitivities.

The prevalence of seemingly contradictory reports underscore the need for a systematic study of relevant factors in a controlled experimental setting, as well as the importance of understanding how antibiotics synergize with other gut perturbations. To close current gaps in understanding, we tracked microbiota dynamics with high temporal and taxonomic resolution during antibiotic treatment in a controlled model system, while isolating variables that influence antibiotic treatment and recovery such as diet, treatment history, and housing co-inhabitants. Our results emphasize that the response of individual bacterial species can be highly affected by the host, the rest of the gut community, and available environmental reservoirs. Our systematic investigation of factors required for resilience and recovery of particular bacterial taxa to antibiotic perturbation lays the foundation for strategies that could mitigate long-term effects of antibiotic treatment on the gut microbiota.

### Microbial load recovers rapidly after massive, transient disturbance from streptomycin

We first sought to determine the extent to which the gut microbiota is disrupted during antibiotic treatment and the timescale over which the community recovers in total abundance and composition. Fifteen germ-free mice distributed across 3 cages were equilibrated with human feces (‘humanized’) from an anonymous donor (Methods) and treated with 20 mg streptomycin per day for five days by gavage (days 0-4). Feces were collected for 16S ribosomal rRNA sequencing immediately before the first gavage, 12 h later (day 0.5), and then every day for 14 days. Three mice were sacrificed on days 0, 1, 5, 8, and 14 for other analyses (Fig. 1A).

**Figure 1:**
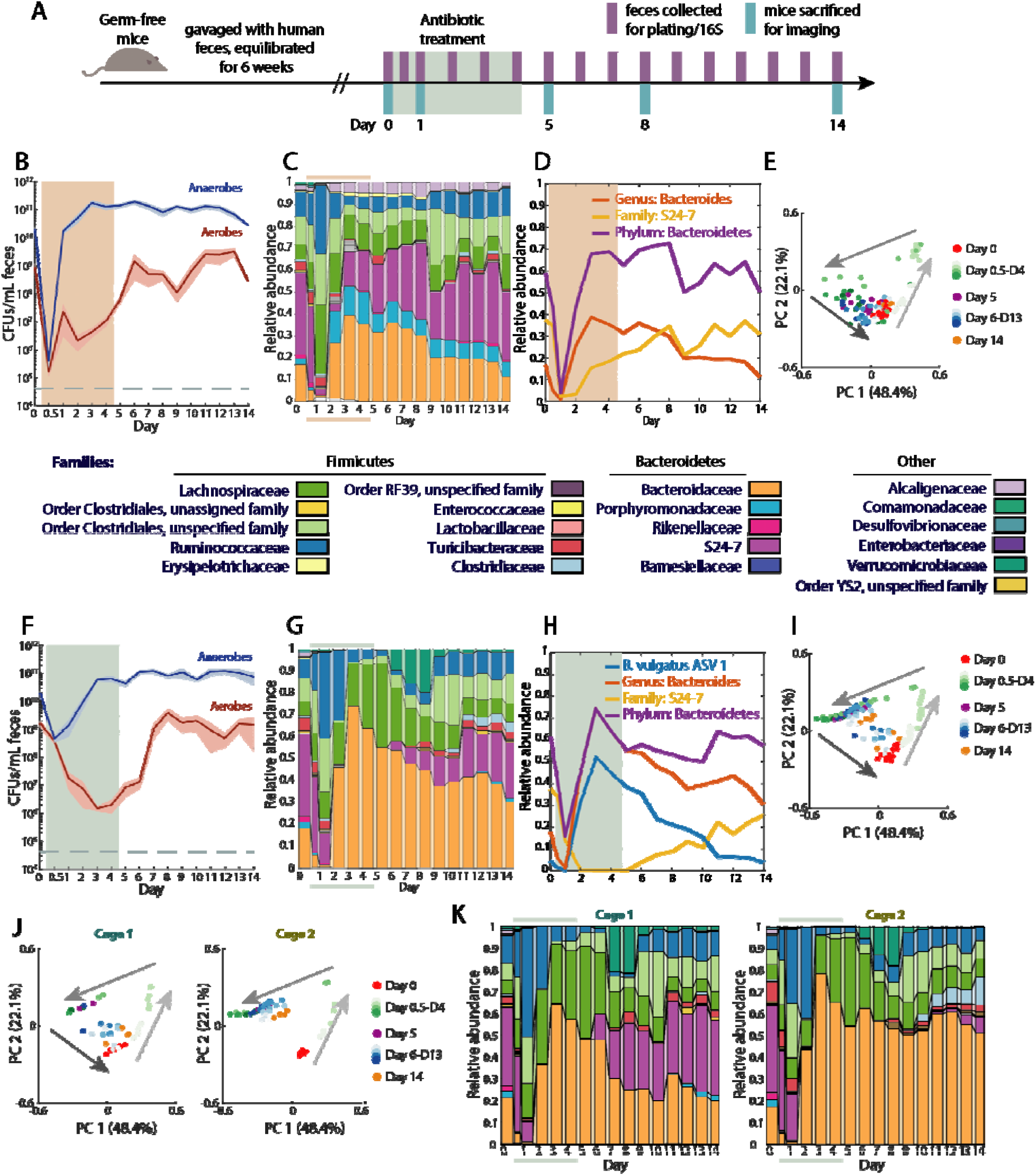
Rapid recovery of bacterial load during antibiotic administration in a human microbiota across antibiotics, with common trajectories of microbiota collapse and recovery but cage-specific end points. (A) Schematic of humanization of germ-free mice, antibiotic treatment, and sampling. Each experiment started with 15 mice, but 3 mice were sacrificed for imaging at each of the designated time points. (B,F) Culturable anaerobic (blue) and aerobic (red) fecal densities in (B) humanized mice treated with 20 mg streptomycin daily for 5 days (*n* = 15), and in (F) humanized mice treated with 3 mg ciprofloxacin twice daily for 5 days (*n* = 14). Bacterial loads recovered during antibiotic treatment (colored area). Error bars: standard error of the mean. (C,G) Family-level community composition in feces in (C) streptomycin-treated humanized mice, and (G) ciprofloxacin-treated humanized mice, (colored bars denote antibiotic treatment period). Humanized mice exhibit similar dynamics under both antibiotics. (D,H) Relative abundance of *Bacteroides*, S24-7 and Bacteroidetes in (D) streptomycin-treated and (H) ciprofloxacin-treated humanized mice. (E,I) Principal coordinates analyses (PCoA) of community composition in humanized mice during (E) streptomycin or (I) ciprofloxacin treatment reveal a conserved trajectory. Analyses used weighted UniFrac distances (Methods). (J) PCoA of humanized mice during ciprofloxacin treatment in two cages uncovers cage-specific differences. (K) Family-level composition in feces from the two cages in (H) shows that S24-7 recovery is stochastic and cage-specific.

We measured the culturable colony forming units (CFUs) per mL in feces in each sample and observed a drop from ∼10^10^ to 10^5^-10^6^ CFUs/mL by day 0.5 (Fig. 1B). Nonetheless, the anaerobic load recovered to >10^10^ CFUs/mL by day 1 and remained thereafter at 10^10^-10^11^ CFUs/mL despite continued drug administration on days 2-4. These rapid dynamics illustrate that even daily sampling frequencies could miss the rapid antibiotic-induced density decrease that occurs after treatment initiates. The density of aerobes was much lower than that of the anaerobes and hence made minimal contribution to the total load, but did not recover until day 6 (Fig. 1B), indicating that treatment continually reconfigured microbiota composition despite the rapid recovery in bacterial load.

### Streptomycin elicits a common trajectory of compositional recovery despite hysteresis and kinetic variation

Using 16S sequencing, we identified amplicon sequence variants (ASVs) and mapped them to taxa using DADA2, which uses error correction to avoid clustering artifacts in operational taxonomic unit (OTU)-based algorithms (Callahan et al., 2016); these ASVs represent predicted ‘species’ in the community. The collapse and recovery of microbiota composition were similar across all mice in multiple cages. Before treatment, mice had similar abundances of Bacteroidetes (comprised of class Bacteroidia) and Firmicutes (classes Bacilli, Clostridia and Erysipelotrichi), with these two phyla accounting for >95% abundance (Fig. 1C). By day 1, Bacteroidetes dropped to <10% (Fig. 1D), with the Firmicutes concomitantly increasing in relative abundance (Fig. 1C). By day 2, the Bacteroidetes returned to pre-antibiotic relative abundances, as did total culturable bacteria (Fig. 1B,D). Betaproteobacteria (represented by the family Alcaligenaceae) increased from ∼2% to ∼5% during treatment and then persisted at that level (Fig. 1C), suggesting that they opportunistically filled a niche opened by the treatment.

These dynamics were evident in a principal coordinates analysis (PCoA), which measures similarity in composition: all mice reached a common point of maximal disturbance along coordinate 1 by day 1 (Fig. 1E). Thus, despite the initial, massive decrease in bacterial load (Fig. 1B), the path to maximum disturbance was deterministic. After day 1, the microbiotas re-equilibrated to the pre-treatment composition along a different path than the one traversed to the point of maximal disturbance (Fig. 1E). In addition to this hysteresis, recovery kinetics varied. Some mice had already returned to the day 0 state by day 5, while others returned more slowly (Fig. 1E), perhaps due to their reliance on re-seeding from other mice in the cage. Nonetheless, all mice followed a similar trajectory in the space of coordinates 1 and 2 (Fig. 1E), suggesting that intermediate states of recovery were similar despite kinetic variation across mice.

### Ciprofloxacin treatment results in similar microbiota dynamics to streptomycin

To determine whether the rapid recovery of bacterial load and phylum-level composition was antibiotic-specific, we treated 14 humanized mice (co-housed in 3 cages) twice daily for 5 days with 3 mg ciprofloxacin, a broad-spectrum DNA gyrase inhibitor. This dosing regimen was chosen based on preliminary experiments in which dosing twice daily had a greater effect than a larger dose once daily (Fig. S1A). The anaerobic compartment of culturable bacteria initially dropped by 100- to 1000-fold by day 1, but recovered to pre-treatment levels by day 3, before administration ended (Fig. 1F), as in streptomycin-treated mice (Fig. 1B). The phylum-level effects of ciprofloxacin treatment on community composition were remarkably similar to those of streptomycin (Fig. 1G). Ciprofloxacin-treated mice traversed a similar trajectory in the space of coordinates 1 and 2 to the point of maximal disturbance as streptomycin-treated mice, despite differences in kinetics (Fig. 1E,I). The similarity wash driven at least in part by the large initial replacement of Bacteroidetes with Firmicutes (Fig. 1C,G,S1B). Bacteroidetes relative abundance dropped to 10-25% by day 1 due to replacement by Firmicutes, but recovered by day 3 and remained stable thereafter (Fig. 1G). This recovery was largely associated with a single ASV matching *Bacteroides vulgatus* (*Bv*) that was at 4% prior to antibiotics but bloomed to >50% in all mice (Fig. 1H), before being replaced by S24-7 family members once treatment ended (Fig. 1H), suggesting a competition between these two families for a common niche (Tropini et al., 2018). We isolated 12 colonies from a mouse sample during *Bv* dominance (day 4) on *Bacteroides*-selective plates; 16S Sanger sequencing revealed that all of the isolates were *Bv*. Whereas the Betaproteobacteria increased in abundance during streptomycin treatment, the Verrucomicrobiae bloomed after ciprofloxacin was removed (Fig. 1G), again indicating opening of favorable niches. Thus, ciprofloxacin and streptomycin influence composition in surprisingly similar manners despite highly distinct molecular effects, although distinctions emerge during recovery.

### Cage-specific effects during recovery from ciprofloxacin indicate importance of environmental microbial reservoirs

After maximal disturbance, one subset of ciprofloxacin-treated mice continued to follow the recovery trajectory of streptomycin-treated mice, while the rest remained similar to the disrupted state (Fig. 1I). Interestingly, these subsets were entirely cage-determined (Fig. 1J,K), suggesting reseeding plays a major role in recovery. This difference was not apparent in class-level taxonomic composition (Fig. S1C). However, an obvious distinguishing feature within the ciprofloxacin-treated mice was the dynamics of the family S24-7, a member of the Bacteroidetes phylum abundant prior to treatment (∼35%) and highly prevalent in homeothermic animals (Ormerod et al., 2016). In one cage that exhibited complete recovery in PCoA space (Fig. 1J), S24-7 disappeared in the latter stages of antibiotic treatment, but re-emerged directly after ciprofloxacin was removed, and re-established at pre-treatment levels (Fig. 1K,S1D). By contrast, in another cage S24-7 was undetectable until day 10, and only recovered to ∼5% by day 14 (Fig. 1K,S1D). In streptomycin-treated mice, S24-7 recovered in all cages, even before cessation of antibiotics (Fig. S1E). A likely explanation for the heterogeneity is that each mouse has a low probability of retaining S24-7 or being recolonized from the cage reservoir, but afterward the rest of the cage shares in S24-7 recovery. These data suggest that gamma diversity of host microbiotas within a cage is an important driver of recovery after antibiotics.

### Antibiotics cause sustained loss in alpha diversity, driven by loss of Bacteroidetes members

Although the culturable densities and phylum-level relative abundances of Bacteroidetes and Firmicutes returned to pre-antibiotic states regardless of microbiota or antibiotic (Fig. 1B,C,F,G), we were interested in long-term effects of perturbation, especially as the loss of S24-7 in some mice (Fig. 1K, S1D) and oligodominance of *Bacteroides* spp. (Fig. 1H) suggested that antibiotic treatment could have long-term effects. To identify one such long-term effect—extinction events—we quantified the alpha diversity, a measure of community resilience (Lozupone et al., 2012). For streptomycin and ciprofloxacin, the alpha diversity of the humanized microbiota dropped during treatment (Fig. 2A,B), with a larger decrease in ciprofloxacin-treated mice despite the smaller decrease in bacterial load (Fig. 1B,F). In both cases, alpha diversity slowly increased after the antibiotic was removed, and re-equilibrated at a level significantly lower than the level pre-treatment (Fig. 2A,B). The diversity increase during recovery could be due to very low-abundance members recovering to detectable levels, or to reseeding from environmental reservoirs.

**Figure 2:**
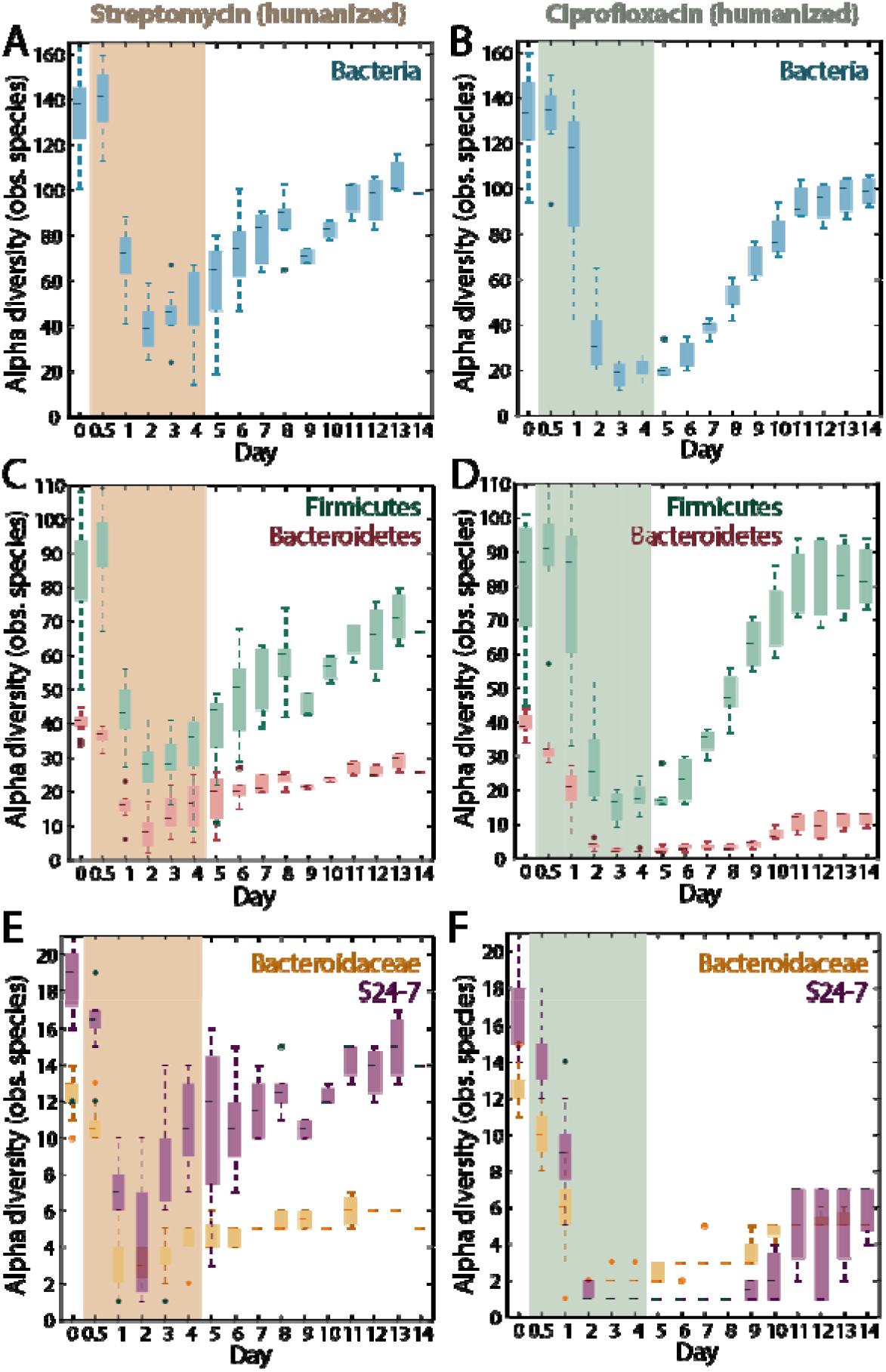
Antibiotic administration alters the state of the microbiota via reductions in the diversity of the Bacteroidetes. (A,B) After treatment, the microbiota stabilizes at lower alpha diversity (observed ASVs; antibiotic treatment period marked in colored area). (C,D) The alpha diversity of the Bacteroidetes (pink) phylum is impacted much more than that of the Firmicutes (green) for all treatments and microbiota. (E,F) The alpha diversity of the S24-7 (purple) and Bacteroidaceae (yellow) families are generally impacted, with S24-7 particularly so by ciprofloxacin.

The sustained decrease was driven largely by taxa within the Bacteroidetes phylum, which permanently decreased in diversity by 36% and 70% in streptomycin and ciprofloxacin-treated humanized mice, respectively (Fig. 2C,D). By contrast, the diversity of Firmicutes was only slightly affected or unaffected (Fig. 2C,D). In streptomycin-treated humanized mice, the decrease in Bacteroidetes diversity was largely accounted for by the Bacteroidaceae family, which includes the *Bacteroides* genus (Fig. 2E); S24-7 family diversity recovered to almost pre-treatment levels (Fig. 2E). By contrast, in ciprofloxacin-treated humanized mice, both Bacteroidaceae and S24-7 diversity experienced a larger decrease than in streptomycin-treated mice (Fig. 2E,F) and remained low after ciprofloxacin was removed. S24-7 diversity decreased to as low as 1 observed ASV, from ∼17 at the start of the experiment, (Fig. 1F), suggesting complete extinction of some species. Faith’s phylogenetic diversity, a complementary measure that incorporates the relatedness of taxa within a community, gave similar conclusions to observed ASVs (Fig. S2A-F). Together, our results reveal that the two dominant phyla exhibit asymmetric vulnerability to two different antibiotics, regardless of *in vitro* sensitivity, with Bacteroidetes loss and recovery influenced by variables such as the initial microbiota (conventional versus humanized) and by cage mates.

### Antibiotic resistance and intra-genus competition during and after ciprofloxacin treatment determines the relative abundances of *Bacteroides* species

The transient decrease in relative abundance and extinction of taxa within the Bacteroidaceae (Fig. 1C,1G,2E,2F) suggests that the Bacteroidaceae experience selection during antibiotic perturbation. To determine whether this behavior was largely driven by the selection for antibiotic resistance, we performed a second antibiotic treatment (Fig. 3A). We employed ciprofloxacin instead of streptomycin because it caused the greater drop in alpha diversity (Fig. 2).

**Figure 3:**
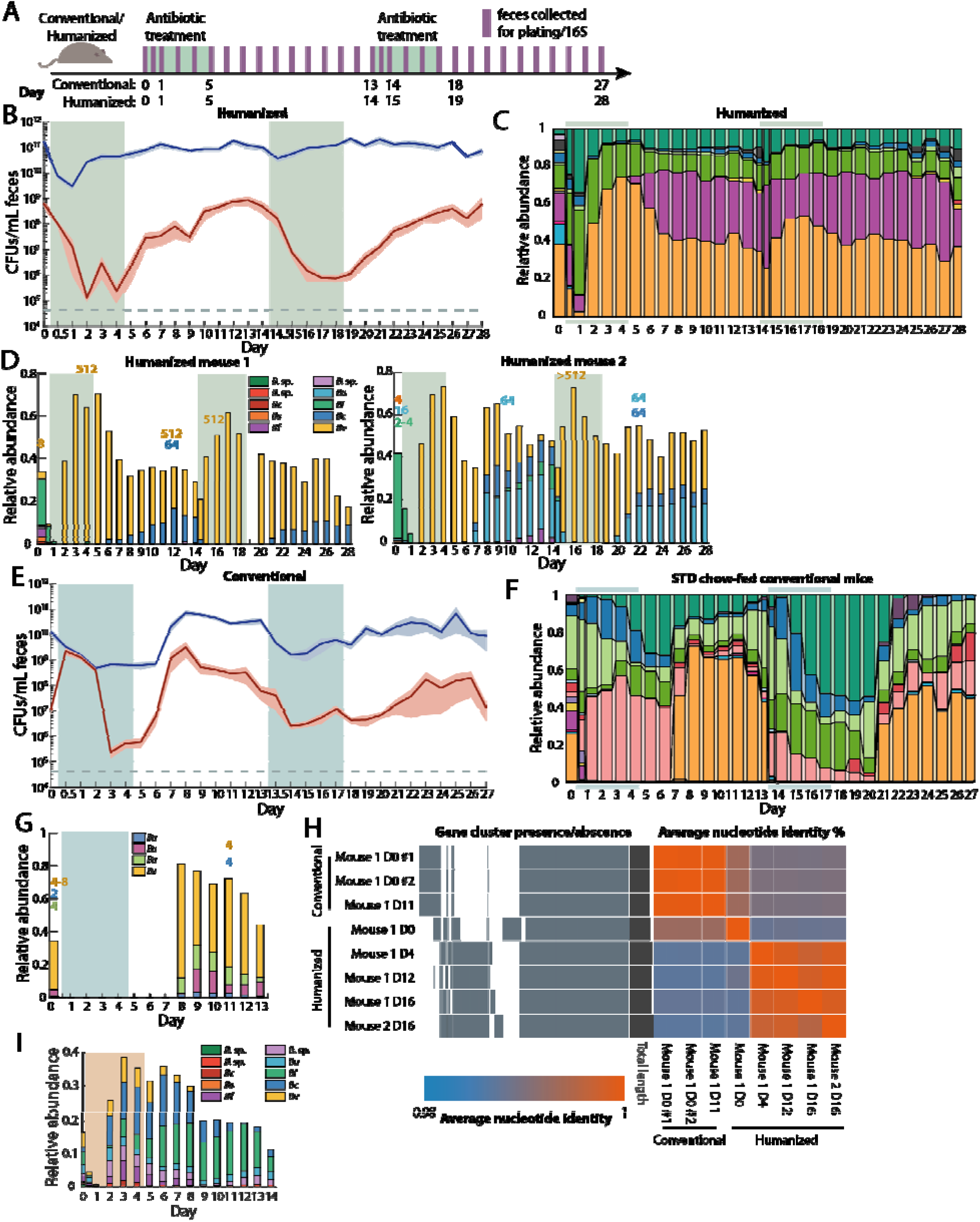
The microbiotas of conventional and humanized mice display distinct signatures under repeated ciprofloxacin treatment. (A) Schematic for treatment of humanized/conventional mice with two courses of ciprofloxacin. (B,E) Culturable anaerobic (blue) and aerobic (red) fecal densities in (B) humanized and (E) conventional mice treated with ciprofloxacin (antibiotic treatment period marked in colored areas) revealed that the effects of the second treatment in conventional mice mirrored those of the first treatment, but humanized mice had a much more robust second response. Error bars: standard error of the mean. (C,F) Family-level community composition in feces of (C) humanized and (F) conventional mice further demonstrated the robustness of humanized mice to the second treatment. (D,G,I) In the presence of differential resistance to antibiotics, resistant strains were dominant during antibiotic treatment but other *Bacteroides* spp. were able to expand after antibiotics. Relative abundance of the 10 most abundant *Bacteroides* ASVs in (D) ciprofloxacin-treated humanized mice, (G) ciprofloxacin-treated conventional mice and (I) streptomycin-treated humanized mice. *B. sp*: unknown *Bacteroides* spp.; *Bc*: *B. caccae*; *Bs*: *B. salyersiae*; *Bf*: *B. fragilis*; *Bu*: *B. uniformis*; *Bi*: *B. intestinalis*; *Bv*: *B. vulgatus*; *Ba*: *B. acidifaciens*. MICs for strains that were isolated are labeled above the day that isolates were derived, colored by the appropriate ASV based on 16S Sanger sequencing. Antibiotic treatment period is denoted by shaded areas. (H) Comparison of gene clusters and average nucleotide identity of *Bacteroides vulgatus* (ASV 1) isolates reveals D0 isolate is different from major isolates from other days, and that the conventional *B. vulgatus* isolates also cluster together.

The anaerobic compartment was negligibly affected by the second treatment (Fig. 3B), consistent with the lack of change in community composition at the family level (Fig. 3C). By day 1 of the second treatment, the class-level composition was almost indistinguishable from the stabilized end of recovery after the first treatment, although this community was different than the baseline community before treatment (Fig. S3A), demonstrating that the initial treatment selected for a more resilient community.

Despite the lack of obvious changes in community composition during the second treatment, throughout both treatments, the Bacteroidaceae dropped to a single, dominant ASV (*B. vulgatus*) and then recovered to 2-5 ASVs after ciprofloxacin was removed (Fig. 3D). This domination during antibiotic treatment was reflected in the alpha diversity, which was negatively impacted (Fig. S3B), indicating a depletion of certain members and filling of niches by other, related species. To examine the extent to which the emergence of ciprofloxacin resistance explains the response of humanized mice to a second treatment, we isolated strains from samples of two mice. We identified strains as *Bacteroides* spp. via 16S sequencing and measured their MICs to ciprofloxacin. The MICs of pre-antibiotic *Bacteroides* isolates (Table S4A) were within the reported range of literature values (2-16 μg/mL) (Goldstein and Citron, 1992). *B. vulgatus* isolates from samples during or after antibiotic treatment displayed increases of >100-fold abundance relative to the isolates from day 0 (Fig. 3D). Whole-genome sequencing of *B. vulgatus* isolates from one mouse revealed that isolates from day 4, 12, or 16, but not the day 0 isolate, possessed a mutation in *gyrA* that confers ciprofloxacin resistance. The genotypes of the day 4, 12 and 16 isolates were highly distinct from those of the day 0 isolate (Fig. 3H), suggesting that the initial population included multiple *B. vulgatus* strains with varying ciprofloxacin sensitivity and that treatment reconfigured their relative abundances. The domination occurred only during antibiotic treatment (Fig. 3D). After antibiotic treatment ceased, although total Bacteroidaceae relative abundance appeared to stabilize, the Bacteroidaceae transitioned rapidly from dominance of a single *B. vulgatus* species on day 6 to multiple *Bacteroides* ASVs (Fig. 3D). This suggests that other *Bacteroides* species are able to reseed and/or recover and then compete effectively with *B. vulgatus* in the absence of antibiotics.

### Conventional and humanized mice have similar recovery trajectories but different kinetics during ciprofloxacin treatment

To determine whether the compositional response of the humanized microbiota was influenced by antibiotic history and/or by the nature of humanization, we carried out a similar double-treatment experiment in conventional mice with a natural laboratory mouse-adapted microbiota that has likely not experienced as much antibiotics as the human donor for the humanized mice. The conventional samples mapped onto approximately the same PCoA trajectories as humanized mice (Fig. S3C), but with slower dynamics; conventional mice on day 8 resembled humanized mice on day 3 (Fig. S3C). Mirroring this delay, the bacterial load (Fig. 3E) and Bacteroidetes abundance (Fig. 3F) did not recover during antibiotic treatment in any mice. Instead, the Bacteroidetes recovered 1 day after treatment cessation, matching the time scale of ciprofloxacin clearance from feces (Fig. S3E), with recovery almost entirely due to increased Bacteroidaceae abundance.

In contrast to a humanized community that exhibited the enrichment and domination of a resistant strain during antibiotic treatment (Fig. 3D), all *Bacteroides* isolates from conventional mice before or after treatment, including the *B. vulgatus* that was the most abundant, had MIC values of 4-8 μg/mL (Fig. 3G); the lack of resistance was consistent with the decreases in bacterial load being similar during the first and second treatments (Fig. 3E). Whole-genome sequencing demonstrated that the day 11 isolate was virtually identical to the day 0 isolate (Fig. 3H), and there were no obvious mutations that would confer antibiotic resistance. Thus, despite identical dosing conditions, the selection for *in vitro* resistance in *B. vulgatus* in a complex community is not pre-destined; the recurrence of a large decrease in bacterial load and similar compositional kinetics at the start of the second treatment suggest that this conclusion extends to the rest of the microbiota as well. Consistent with the lack of selection of a resistant *Bacteroides* species during antibiotics, there was no domination by a single species, and multiple *Bacteroides* spp. were maintained in the period after antibiotic treatment.

Concomitant with the sensitive response of the *Bacteroides*, we observed that S24-7 was highly responsive (similar to its decrease during antibiotic treatment of humanized mice); however, in conventional mice, S24-7 never recovered to the high densities seen in some humanized mice (Fig. 3C,F). In sum, our data suggest that the conventional mouse microbiota is more sensitive to treatment than the humanized mouse microbiota. The effects of the second treatment were remarkably similar to those of the first treatment: ∼10-100-fold drop in CFUs/mL (Fig. 3E), and the Bacteroidetes were replaced by a combination of Firmicutes and Verrucomicrobiae (Fig. 3F). The alpha diversity failed to recover to the same level as before the second treatment (Fig. S3D), suggesting further elimination of community members. Bacteroidetes species were undetectable during treatment and remained so until 3 days after treatment stopped (Fig. 3F), and S24-7 was driven to undetectable levels in all mice immediately upon the second treatment (Fig. 3F). Thus, an inability to recover completely from the first treatment predisposes the conventional microbiota to a more profound disturbance upon the second treatment.

### *In vitro* resistance of the *Bacteroides* genus contrasts with its *in vivo* response to streptomycin

Antibiotic resistance appears to explain the response of *B. vulgatus* to ciprofloxacin, hence we wondered whether the recovery of the *Bacteroides* after day 1 of streptomycin treatment (Fig. 1D) was also due to the emergence of resistance. However, previous studies suggested that anaerobic bacteria are resistant to aminoglycosides such as streptomycin *in vitro* (Bryan et al., 1979), making it surprising that the Bacteroidetes decreased dramatically in abundance during the first 12 h of streptomycin treatment (Fig. 1C). We measured streptomycin MICs *in vitro* of *Bacteroides* laboratory strains and strains isolated from the pre-treatment donor sample (SI). The majority exhibited no adverse growth effects at 512 μg/mL streptomycin, the highest dose tested (Fig. S4A), supporting the resistance of the *Bacteroides* genus. Despite being present at 17% before treatment, surprisingly they experienced a similarly massive (∼10^5^-fold) initial decrease in total abundance as the rest of the community upon treatment, given that they are straightforwardly cultured on BHI plates. To test whether the drop in *Bacteroides* at the start of treatment arose indirectly from the killing of other species, affecting cross-feeding and nutrient availability, or from host-related effects, we monocolonized germ-free mice with *B. thetaiotaomicron* and treated with streptomycin. CFUs dropped during the first 24 hours to a similar extent as in humanized mice (Fig. S4B), indicating that the host somehow reversed the natural resistance of *B. thetaiotaomicron* to streptomycin. These data demonstrate that *in vitro* susceptibility measurements are not necessarily predictive of *in vivo* response.

Additionally, we did not observe dominance of *B. vulgatus* (or any other *Bacteroides* spp.) in streptomycin-treated humanized mice, in contrast to ciprofloxacin-treated humanized mice (Fig. 3I), supporting the hypothesis that dominance arises out of selection of resistant strains during antibiotic treatment. In conventional mice, dominance was not observed (Fig. 3G) because there did not appear to be antibiotic resistant strains present before antibiotic treatment that could take advantage of the sensitivity of related strains and outcompete them during antibiotics. In the case of streptomycin, the lack of selection is likely due to the near-universal resistance across the *Bacteroides* genus.

### Antibiotic treatment increases luminal mucus

Given that domination of ciprofloxacin-resistant *Bacteroides* spp. occurred only during antibiotic treatment but was disrupted by the expansion of non-resistant strains after cessation of antibiotics, we wondered whether factors involving the host environment, namely nutrient availability, affected taxonomic dynamics. Since the Bacteroidetes recover during antibiotic treatment, and many *Bacteroides* spp. are known mucus utilizers (Salyers et al., 1977; Sonnenburg et al., 2005), we hypothesized that mucus availability might increase during antibiotic treatment. We fixed the tissues of humanized mice sacrificed on days 0, 1, 5, 8, and 14 and mounted them for DNA and mucus staining and imaging as previously described (Earle et al., 2015; Johansson and Hansson, 2012) (Fig. 4A,B).

**Figure 4:**
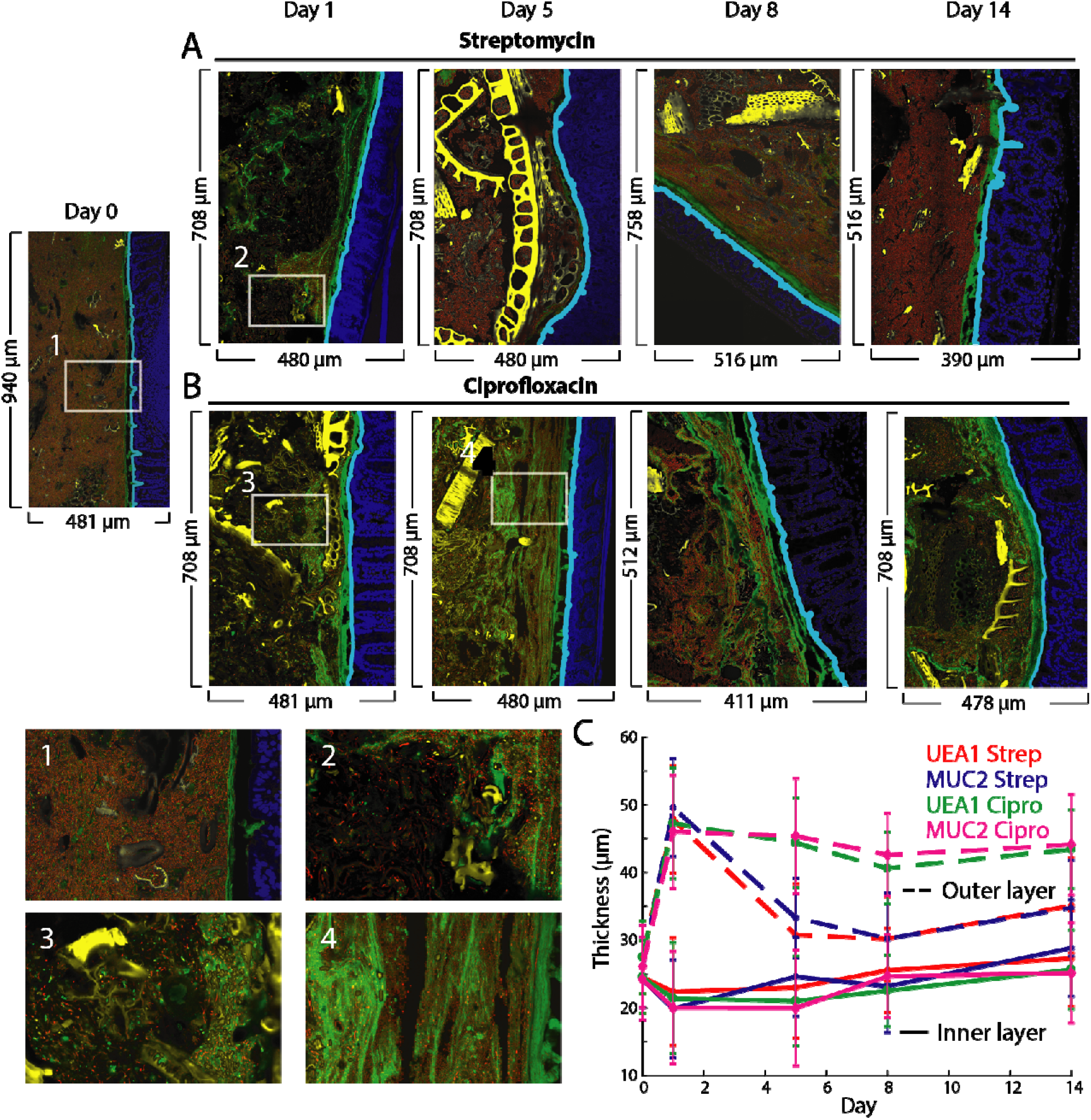
Antibiotic treatment increases loose intestinal mucus. (A,B) Imaging of the distal colon of humanized mice before, during, and after treatment with (A) streptomycin and (B) ciprofloxacin reveals a sustained increase in loose luminal mucus. Sections were stained with DAPI (epithelial DAPI, blue; luminal DAPI, red) and MUC2 (green). Computationally subtracted debris appears in yellow. Zoomed-in panels: (1) Dense community before antibiotics with little luminal mucus and a tight mucus layer. (2,3) Massive decrease in luminal DAPI signal 1 day after streptomycin and ciprofloxacin treatment. (4) Recovery of luminal DAPI but persistence of loose luminal mucus. (C) Thickness of the tight mucus (“inner layer”) remains approximately constant during treatment, while the looser luminal mucus (“outer layer”) proliferates during treatment with both antibiotics and persists after ciprofloxacin treatment.

We utilized the software *BacSpace* (Earle et al., 2015) to identify the epithelial boundary, to segment bacteria from plant material and other debris in the lumen, and to quantify the thickness of the mucus layer. For both antibiotics, the thickness of the tight mucus layer was relatively constant over time (Fig. 4C). However, there was an obvious expansion of Muc2 signal into the lumen upon treatment (Fig. 4C), similar to the increase in mucus in germ-free mice (Johansson et al., 2008).

In ciprofloxacin-treated humanized mice, this increase in the loose mucus layer persisted throughout treatment and recovery, suggesting that the increase was not due to direct effects of the ciprofloxacin on the mucus since ciprofloxacin is cleared by day 7 (Fig. S3E), while in streptomycin-treated mice, the loose mucus returned to baseline by day 5 (Fig. 4C). The Verrucomicrobiae phylum, which includes the mucin-degrading *Akkermansia muciniphila*, did not noticeably change in abundance during or after streptomycin treatment (Fig. 1C), but bloomed after ciprofloxacin treatment was removed (Fig. 1G). Additionally, whole genome sequencing revealed that *Bacteroides* strains isolated from post-ciprofloxacin fecal samples possessed more mucin and host carbohydrate-degrading genetic modules than strains isolated before antibiotic treatment (Fig. S4C), suggesting that the capability to eat loosened mucus may drive the expansion of *Bacteroides* spp. and other mucus-responsive phyla (Fig. 1C,G) and counteract the domination of *B. vulgatus* (Fig. 1H,3D).

### A polysaccharide-deficient diet disrupts recovery after ciprofloxacin treatment

Previous research has demonstrated that feeding mice a diet lacking MACs thins the tight mucus layer (Desai et al., 2016; Earle et al., 2015); bacteria deprived of dietary carbohydrates must utilize mucin-derived polysaccharides instead. A MAC-deficient diet also selects for mucin utilizers like *A. muciniphila* (Earle et al., 2015; Marcobal et al., 2013), which may compete with *Bacteroides* spp. for mucus carbohydrates. We hypothesized therefore that the reduction in mucus during a MAC-deficient diet would antagonize the proliferation of loose mucus during antibiotic treatment (Fig. 4C) and thereby hamper the recovery of mucin utilizers such as the Bacteroidaceae.

We transitioned humanized mice (with the same microbiota as in Fig. 1–3) onto a MAC-deficient diet for two weeks, and then began ciprofloxacin treatment (Fig. 5A-D). As predicted, the Bacteroidaceae did not completely recover until after cessation of antibiotics in MAC-deficient diet-fed (MD) mice, unlike standard diet-fed (STD) mice (Fig. 5D,1G). This delayed recovery was accompanied by incomplete recovery of other families within the Bacteroidetes; the S24-7 family was completely eliminated from MD mice (Fig. 5D). Moreover, the alpha diversity of MD mice started somewhat lower than mice that continued on the standard diet (Fig. 5C, *p* = 0.02), and dropped even more with antibiotic treatment (∼6-fold in MD mice vs. ∼4-fold in STD mice; Fig. 5C), indicating that the antibiotic insult acted in an additive fashion with diet shifts. In MD mice, the anaerobic (and total) bacterial load (Fig. 5B) exhibited similar dynamics to mice on a standard diet (Fig. 1F), although the aerobic compartment decreased more and recovery was slower than in STD mice (Fig. 5B, 1F), mirroring recovery delays in alpha diversity (Fig. 5C) and community composition (Fig. 5D).

**Figure 5:**
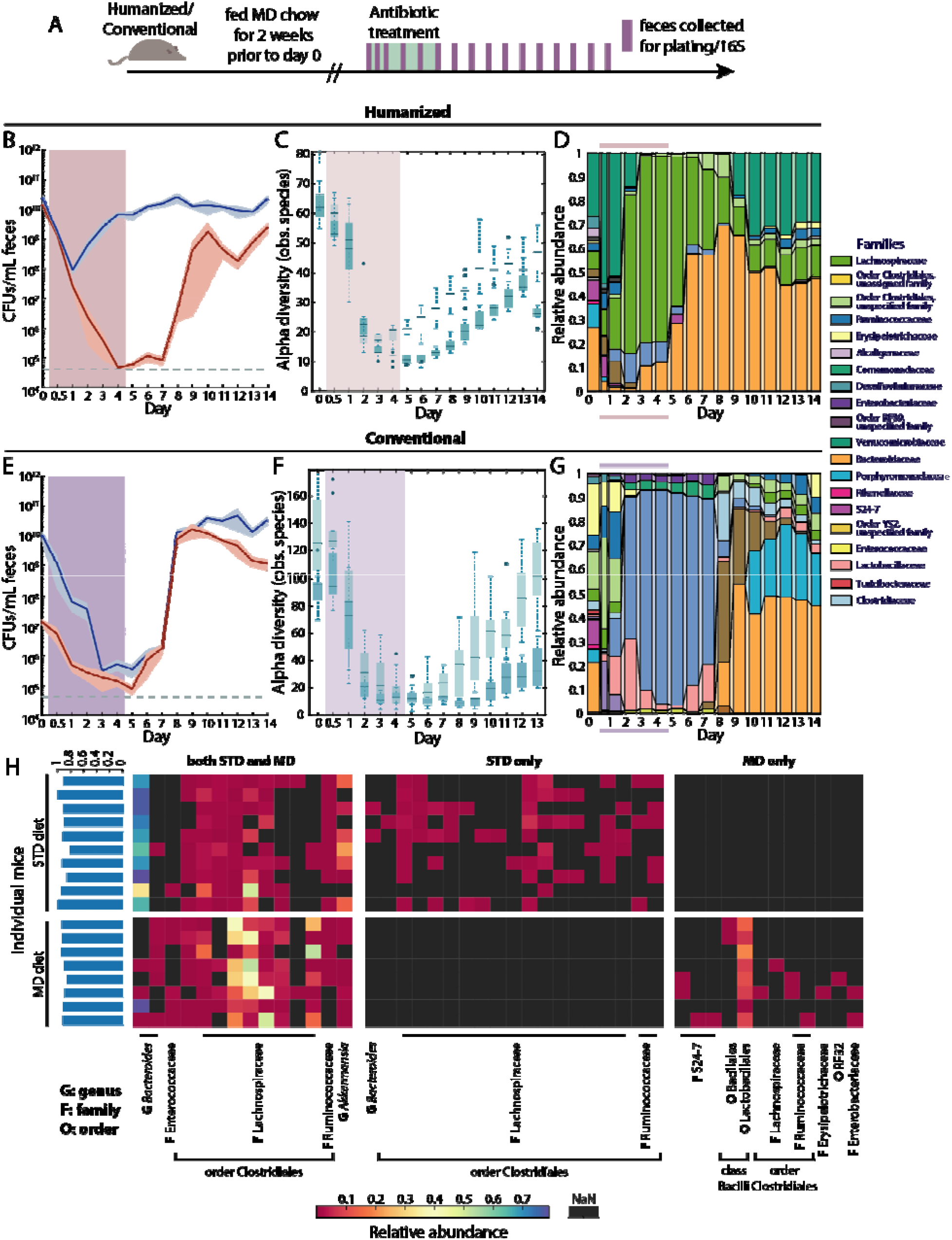
A MAC-deficient diet sensitizes the microbiota to ciprofloxacin. (A) Schematic for antibiotic treatment and sample collection of humanized/conventional mice fed with STD or MD chow. (B,E) Culturable anaerobic (blue) and aerobic (red) fecal densities in (B) humanized or (E) conventional mice fed MD chow revealed that the aerobic compartment is sensitized to ciprofloxacin in humanized mice, and both compartments are sensitized in conventional mice. Error bars: standard error of the mean; antibiotic treatment period marked in colored area. (C,F) Alpha diversity decreases more in MD mice (dark blue) than in STD mice (light blue) after ciprofloxacin treatment. (D,G) The family-level composition in feces of MD mice substantially differs from STD mice (Fig. 1F,K,L). (H) Heatmap of the core microbiome at the nadir of alpha diversity in STD and MD humanized mice. Shared ASVs are clustered on the left and STD- and MD-specific ASVs appear on the right. The percentage of the total relative abundance that is accounted for by the shared core microbiome is ∼90% across almost all mice (blue bars, left).

To determine whether the effects of the diet shift were microbiota-dependent, we carried out a similar experiment with conventional mice (Fig. 5A). In these mice, the bacterial load was dramatically affected by ciprofloxacin treatment, with a >10,000-fold decrease in anaerobic density (Fig. 5E), in contrast to the ∼10- to 100-fold drop in conventional STD mice (Fig. 3E). Similar to humanized mice (Fig. 5B-D), recovery was also delayed in MD versus STD conventional mice (Fig. 5E-G). The decrease in alpha diversity was even more prolonged in MD conventional mice than in MD humanized mice, despite a higher starting diversity (Fig. 5F). Taken together, these data indicate that the shift to a MAC-deficient diet exacerbates the effects of ciprofloxacin, particularly for the more sensitive conventional mouse microbiota.

### Ciprofloxacin selects for a common core of species in humanized mice regardless of diet

Given the decrease in diversity after ciprofloxacin treatment due to a diet switch, we wondered if species extinction exhibited increased stochasticity in MD compared with STD mice. Thus, we focused on the species present at the time of minimum alpha diversity in humanized mice (Fig. 5H). Some species showed diet-specific sensitivity: 6 ASVs found in at least 50% of STD mice and 3 ASVs found in >50% of MD mice were not present in the other diet (Fig. 5H). Nonetheless, a core of 14 ASVs was present at the time of lowest diversity in at least one MD and one STD mouse, with 6-11 of the core ASVs present in each STD mice and 8-12 present in each MD mice (Fig. 5H). Four of the 16 core ASVs were detected in all STD mice, and five were present in all MD mice (Fig. 5H). Overall, this core set of ASVs represented ∼92% of the total abundance in STD mice and ∼90% in MD mice (Fig. 5H). Thus, ciprofloxacin selects for a specific cohort of microbes at the point of maximal disturbance regardless of diet, even with a larger decrease in load and diversity in MD compared with STD mice.

### An initial antibiotic treatment can condition the *Bacteroides* response of a human microbiota to a second treatment

To determine whether the response of the *Bacteroides* and other taxa to ciprofloxacin depends on the overall composition of the microbiota, we humanized mice for 6 weeks with a sample from a different donor. The microbiota of mice from this second donor (H2) had a higher fraction of Bacteroidetes and a lower fraction of Firmicutes than the first donor (H1) (Fig. 6A,B). We treated these H2 mice with ciprofloxacin as previously. As in all previous experiments, the relative abundance of the Bacteroidetes dropped during the first day with a concurrent bloom of Verrucomicrobiae (Fig. 6B). However, the response thereafter was dramatically different than in H1 mice. Instead of recovery of the Bacteroidetes during days 2-5, the Lachnospiraceae bloomed to >70% on day 3 (Fig. 6B). On day 4, the Barnesiellaceae, a sister family of the Bacteroidaceae and S24-7, began to recover (Fig. 6B). S24-7 did not begin recovery until day 6, and the Bacteroidaeceae were undetectable until day 9 (Fig. 6B), similar to H1 mice after a MAC-deficient diet shift (Fig. 5D), suggesting that the H2 microbiota is sensitized to ciprofloxacin treatment.

**Figure 6:**
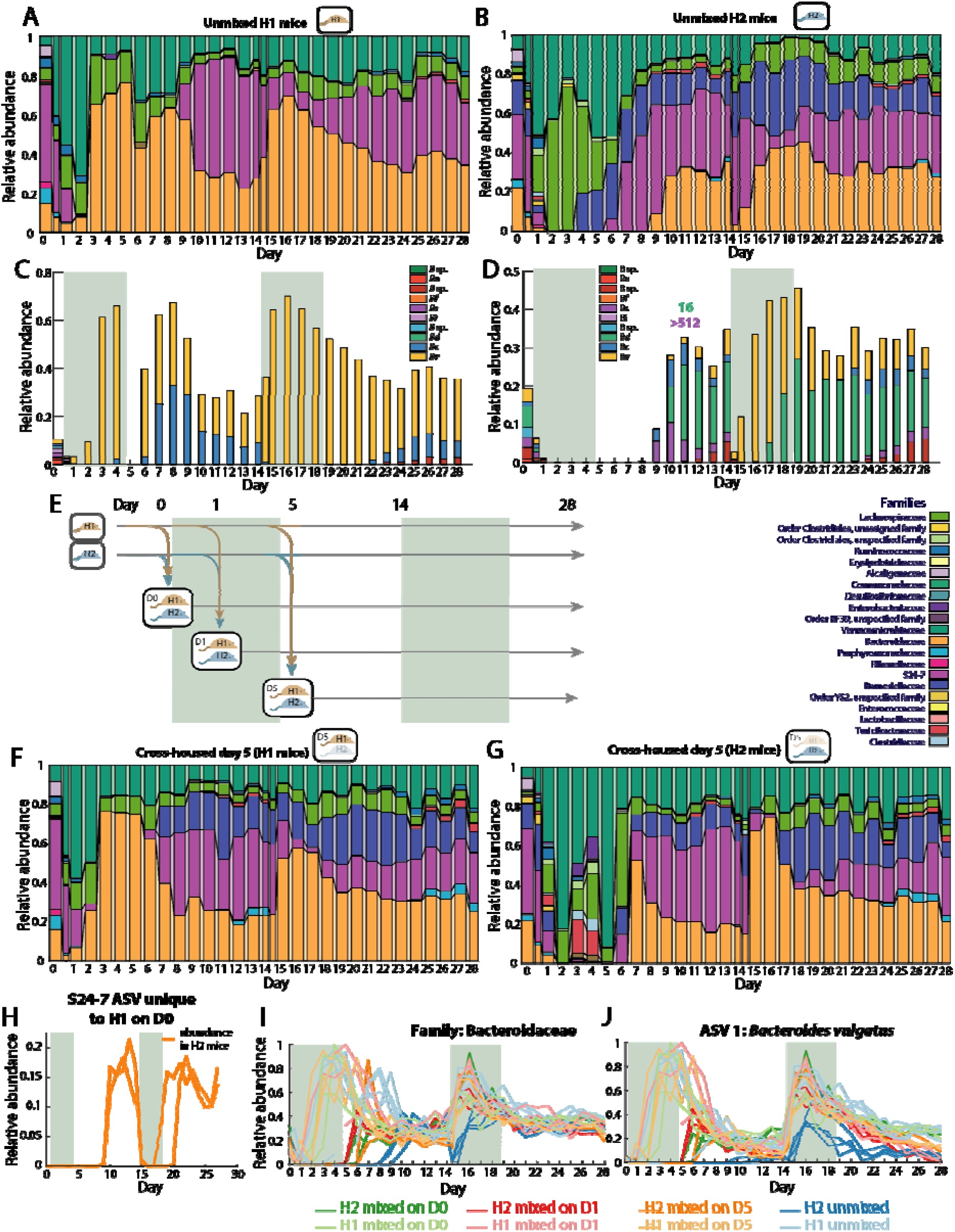
Cross-housing-mediated invasion of *Bacteroides* improves microbiota resilience. (A,B) Family-level community composition in feces of mice colonized with donor samples 1 or 2 (H1 (A, *n*=5) or H2 (B, *n*=4), respectively) during two ciprofloxacin treatments revealed delayed recovery of the Bacteroidetes in H2 mice during the first treatment but not the second. (C,D) Patterns of *Bacteroides* recovery differed between (C) H1 and (D) H2 mice, as revealed by the relative abundances of the 10 most abundant *Bacteroides* ASVs (abbreviations same as in Fig. 5H,I). MICs were calculated for two isolates from H2 mouse samples (labeled above the day in which the isolates were derived). Antibiotic treatment periods are denoted by colored areas. (E) Schematic for cross-housing of mice with different microbiotas at various timepoints before, during, and ciprofloxacin treatment. (F,G) Family-level community composition in feces in (F) H1 mice mixed with H2 mice on day 5 (*n*=3) and (H) H2 mice mixed with H1 mice on day 5 (*n*=2). Cross-housing stimulates Barnesiellaceae blooming after the first treatment in H1 mice, and resilience of the *Bacteroides* during the second treatment in H2 mice (as in unmixed H1 mice). (H) Relative abundance of a S24-7 ASV that was only present in H1 mice on day 0 but then bloomed in H2 mice, highlighting a potential seeding event. (I,J) Relative abundances of (I) Bacteroidaceae and (J) the dominant *B. vulgatus* ASV in all H1 and H2 mice, including cross-housed and unmixed mice, during a double course of ciprofloxacin treatment. The *B. vulgatus* ASV becomes dominant in all H1 mice during both treatments, but only during the second treatment in H2 mice. The *B. vulgatus* abundance increases more rapidly and reaches a higher level in cross-housed H2 mice compared with unmixed H2 mice.

The *B. vulgatus* ASV that was dominant in H1 mice throughout the two antibiotic treatments (Fig. 6C) was initially present at similar levels in H2 mice (∼3.5% in H2 vs. 1.8% in H1), but did not recover until day 10, and then only to ∼6.5% by day 14 (Fig. 6D), unlike its rapid recovery during treatment in H1 mice (Fig. 6C). A second ciprofloxacin treatment resulted in a Bacteroidaceae response similar to that of H1 mice during a first treatment: Bacteroidaeceae relative abundance dropped rapidly but then recovered to levels comparable to H1 mice by the second day (day 16) (Fig. 6B). The *B. vulgatus* ASV dominated throughout this second recovery (Fig. 6D). Notably, we also isolated a *B. salyersiae* from day 11 samples of H2 mice with MIC >512 μg/mL that decreased to almost undetectable levels during the second treatment (Fig. 6D), indicating that *in vitro* sensitivity is only partly predictive of the response *in vivo*, much like the initial drop in the *Bacteroides* during streptomycin treatment despite its antibiotic resistance. These observations suggest that there is an ecological landscape with distinct Bacteroidaeceae recovery behaviors, and that it is possible for a microbiota to move between them through antibiotic treatment.

### Invasion during cross-housing accelerates recovery from ciprofloxacin

The large dynamics in bacterial load and taxonomic composition during ciprofloxacin treatment suggested the potential for invasion from environmental reservoirs, including pathogen colonization. Anticipating the microbiota-specific responses of individual taxa to ciprofloxacin treatment that we identified between H1 and H2 mice (Fig. 6A,B), we also co-housed subsets of H1 and H2 mice (henceforth referred to as cross-housed) on days 0, 1, and 5 (Fig. 6E). These three time points were selected to evaluate whether a particular window of time exists during which invasion is facilitated. Mice underwent two ciprofloxacin treatments, as before. With the exception of the transfer and blooming of the family Barnesiellaceae, which was not present in H1 mice, the compositional dynamics of H1 mice were relatively unaffected by cross-housing with H2 mice (Fig. 6F). By contrast, recovery of the Bacteroidaceae in H2 mice was accelerated by cross-housing (Fig. 6G,I, S5A,B). Cross-housing on day 5 led to multiple waves of transient blooming involving families that were not major components of the stabilized community (Fig. 6G). Moreover, an S24-7 ASV that was initially absent from all H2 mice invaded to ∼15% abundance from H1 mice after cross-housing (Fig. 6H), suggesting that survival of S24-7 species is dependent on reseeding.

Interestingly, by day 10, the family-level composition of all H1 and H2 mice was similar regardless of cross-housing (Fig. 6A,B,F,G). However, the decrease in Bacteroidaceae relative abundance in H2 mice upon the second antibiotic treatment that occurred in non-cross-housed mice (Fig. 6B,I) was essentially eliminated by cross-housing (Fig. 6G,I, S5A,B). In all cross-housed H2 mice, the *B. vulgatus* ASV that was dominant during treatment in H1 mice bloomed to 20-30% directly after the first treatment (Fig. 6J), similar to the levels in H1 mice and consistent with the transfer of the resistant strain from H1 mice. After the first treatment, the dynamics of this ASV in cross-housed H2 mice were essentially the same as in H1 mice, with a quantitatively similar increase in abundance upon start of the second treatment (Fig. 6J); in non-cross-housed H2 mice, the relative abundance of this ASV initially decreased despite being similar to cross-housed H2 mice at the start of the second treatment (Fig. 6J), suggesting that both community context and antibiotic resistance play a role in *Bv* dynamics and dominance. Thus, cross-housing effectively shifts the H2 microbiota toward the response of the H1 microbiota to ciprofloxacin, indicating that environmental reservoirs can reprogram the microbiota toward more robust antibiotic recovery of certain species.

### Reduction of environmental reservoirs impairs microbiota recovery and increases stochasticity

The observation of invasion of taxa from cagemates (Fig. 6F,H) indicates that microbes from other hosts are an important resource for community recovery after antibiotics. Thus, we hypothesized that singly housing mice to mimic the increased sanitation prevalent in Western society would negatively impact recovery. We compared the effects of streptomycin treatment on co-housed and singly housed mice; streptomycin was selected over ciprofloxacin based on the larger drop in bacterial load (Fig. 1B,F), and hence we hypothesized that stochastic extinction would be more likely to play a role. We selected 6 mice at random from a cohort of 11 co-housed ex-germ-free, conventionalized mice and separately housed them in their own cages (Fig. 7A). In all mice, the anaerobic load decreased by 10^2^-10^5^ over the first 24 h and the aerobic load decreased by ∼10^3^ (Fig. 7B). However, the rapid recovery thereafter in co-housed mice was absent in singly housed mice (Fig. 7B), indicating the absence of species that could flourish in the presence of streptomycin.

**Figure 7:**
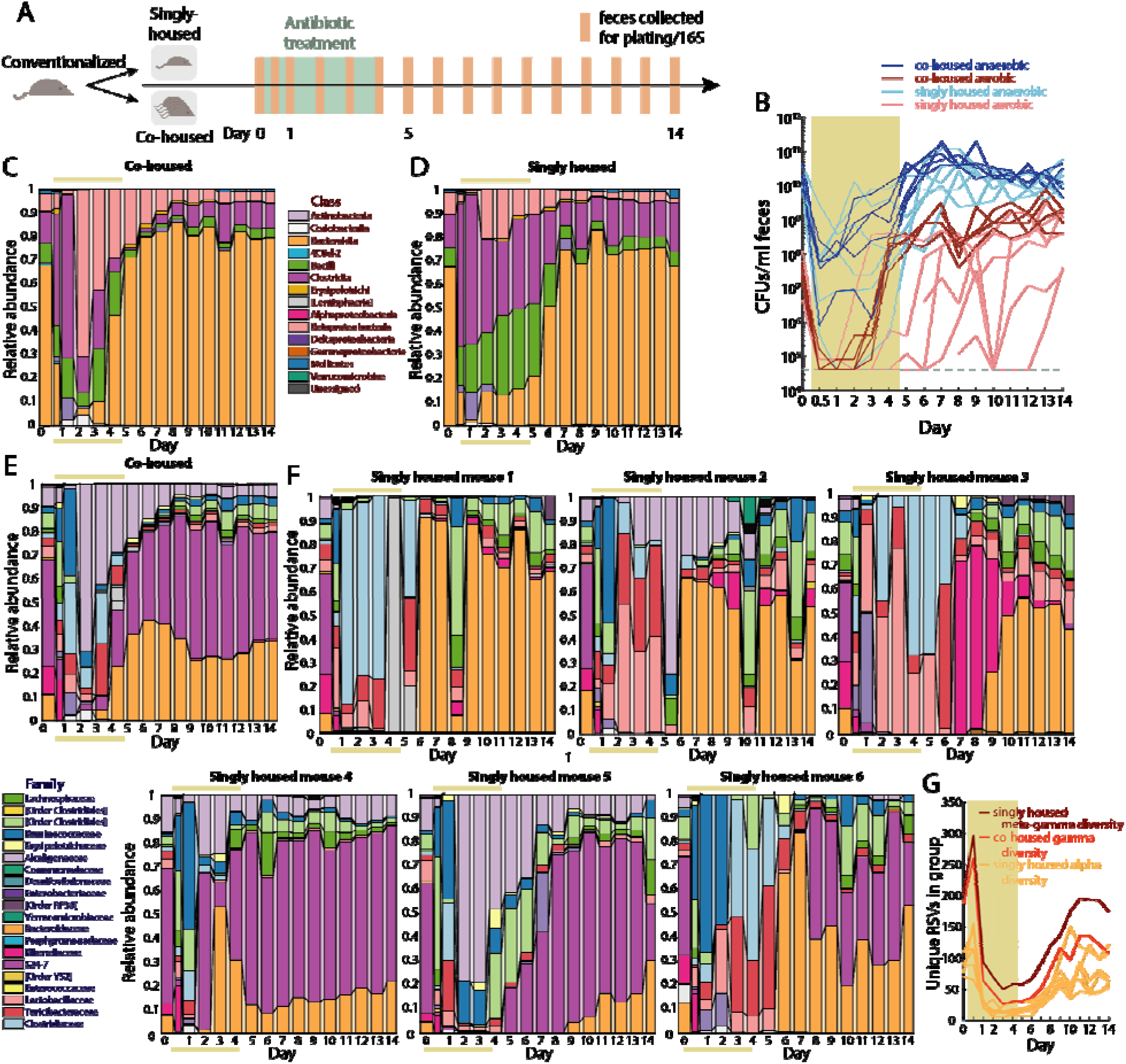
Co-housing enables robust recovery after streptomycin treatment. (A) Schematic for antibiotic treatment and sample collection during housing experiment. (B) Culturing of the anaerobic (blue) and aerobic (red) fecal densities in co-housed (dark lines) and singly housed (light lines) streptomycin-treated conventionalized mice (colored area denotes antibiotic treatment period) revealed delayed recovery of the aerobic compartment. (C,D) Class-level community composition in feces in (C) co-housed and (D) singly housed mice revealed delayed recovery of the Bacteroidetes phylum in singly housed mice. (E) Family-level community composition in feces in co-housed mice (*n*=5) shows robust recovery of the Bacteroidaeceae during streptomycin treatment. (F) Family-level community composition in feces reflects the heterogeneous recovery pattern across six singly housed mice. (G) The noisiness of reseeding in singly housed mice without a common environmental reservoir was evident from the gamma diversity (unique ASVs) of co-housed mice, alpha diversity of singly housed mice, and meta-gamma diversity of singly housed mice treated as if they were a group.

The co-housed conventionalized mice showed similar compositional dynamics (Fig. 7C,E) to our previous co-housed, streptomycin-treated humanized mouse experiment (Fig. 1C), indicating that humanized and conventional microbiotas have similar responses to streptomycin. By contrast, the singly housed mice exhibited dramatically different compositions from co-housed mice (Fig. 7C-E) and from each other (Fig. 7F); no two mice were even qualitatively similar during treatment and recovery. While the Bacteroidaceae recovered during treatment in co-housed mice (Fig. 7E, S6A), they did not recover in many singly housed mice until after antibiotics were removed (Fig. 7F,S6A). S24-7 recovered in all co-housed mice (Fig. 7E,S6B) but in only half of the singly housed mice (Fig. 7F,S6B); in some cases singly housed mice experienced transient blooms of families that were at low abundance or absent in co-housed mice.

After the cessation of streptomycin treatment, the individual diversity of 5 of the 6 singly housed mice was significantly lower than the gamma diversity of the co-housed mice (Fig. 7G). We hypothesized that the slower and more variable recovery dynamics of singly housed mice was a result of stochastic extinctions with no reseeding reservoir due to the absence of cagemates. We therefore predicted that we would observe different cohorts of species across the individual mice after treatment. To test this hypothesis, we computed the total number of unique ASVs represented by the singly housed mice, akin to gamma diversity had the mice been co-housed. As hypothesized, this “meta”-gamma diversity was higher than the gamma diversity of the co-housed mice (Fig. 7G). Taken together, our findings reinforce the importance of environmental reservoirs for robust recovery of the microbiota, and also highlight that antibiotic-treated microbiotas are susceptible to colonization by important commensals as well as invasion by pathogens.

## Discussion

Our study provides important insight into the response to antibiotics of individual strains, taxa, and the entire community. The death of virtually all *Bacteroides* cells in streptomycin-treated mice (Fig. 1B,C) despite their resistance *in vitro* (Fig. S4) illustrates that the *in vitro* sensitivity of commensals is not necessarily predictive of their response to antibiotics when living within the context of a complex ecosystem housed in a mammalian host. The death of *Bacteroides* spp. due to streptomycin treatment in all colonization conditions tested (humanization (Fig. 1B,C), conventionalization(Fig. 7B-D), monocolonization with *B. thetaiotaomicron* (Fig. S4B)) suggests that the drug may be disrupting the host (e.g. through charge-mediated mucosal damage), and the *Bacteroides* dynamics may be the consequences of that disruption and the subsequent repair. Tissue imaging revealed increases in mucus thickness due to both streptomycin and ciprofloxacin treatment (Fig. 4) that could dictate community dynamics, motivating future studies of gut biogeography during perturbations. We uncovered several common features of the response to antibiotic treatment across microbiotas and between streptomycin and ciprofloxacin. One of the most striking commonalities was the recovery of bacterial load during antibiotic treatment, an effect that was common to all five antibiotic treatments tested in humanized mice (Fig. S7A,B). Thus, despite the lack of predictability based on the behavior of isolated species, community features such as density may have predictable responses to antibiotics independent of mechanism of action.

While the dynamics of many taxa are potential fodder for future investigation, the Bacteroidetes phylum, and the S24-7 family and *Bacteroides* genus in particular, were of particular interest throughout our study due to notable dynamics. In all five antibiotics in our pilot, Bacteroidetes relative abundance was either maintained throughout, or in the case of the transcriptional inhibitor rifaximin showed dynamics during treatment that was similar to streptomycin and ciprofloxacin (Fig. 1C,F, S7B). Dominance of a single *Bacteroides* species was observed in both ciprofloxacin (*B. vulgatus*, Fig. 1H), as well as in clindamycin (*B. ovatus*, Fig. S7C) in which the *Bacteroides* had already increased in abundance by day 1. Based on our ciprofloxacin analysis, we surmise that the domination of the *B. ovatus* strain likely results from intrinsic resistance to clindamycin, as has been found at high frequency in surveys of clinical isolates (Karlowsky et al., 2012). Nonetheless, the elimination of a resistant *B. salyersiae* strain from our mice during ciprofloxacin treatment (Fig. 6D), and perhaps other resistant taxa, motivates future studies of the interactions among antibiotic resistant strains. Interestingly, the lack of domination of a single *Bacteroides* strain when there is pan-resistance (streptomycin) or likely pan-susceptibility (rifaximin) (Finegold et al., 2009) suggests that coexistence is maintained during a perturbation as long as there is a level playing field. Just as we identified a core set of species that survived ciprofloxacin treatment regardless of host diet (Fig. 5H), there was a distinct core of 24 ASVs that appears in at least 5 of the 10 mice in our pilot study that accounts for 70.4±13.1% of the microbiota on day 3 of treatment, with 88.6±6.0% of that accounted for by *Bacteroides* or S24-7 species (Fig. S7D). These data suggest that a particular set of interspecies interactions establish resilience to perturbations in general, and that focus on S24-7 and *Bacteroides* spp. is likely warranted for other antibiotics as well, although it may also be the case that a non-*Bacteroides* resistant species or taxon can also achieve domination in other microbiotas.

Our results indicate that caution must be taken when designing antibiotic experiments. In cases such as streptomycin, it was critical to sample at high frequency to capture the large changes in bacterial load and composition (Fig. 1B,C). Strikingly, sampling once daily would have led to the erroneous conclusion that streptomycin has no effect on bacterial load, rather than the 10^5^-fold decrease that we observed after 12 h (Fig. 1B); such effects may explain differences between published reports (Ng et al., 2013; Stecher et al., 2007; Thompson et al., 2015). In a pilot experiment, we detected little to no reduction in CFUs/mL in humanized mice during daily sampling throughout metronidazole treatment (Fig. S7A), and very little effects on microbiota composition aside from blooming of the Verrucomicrobiae (Fig. S7B). More rapid sampling may thus be necessary to quantify the rapid effects of metronidazole on the microbiota, which is remarkably resilient to this perturbation (Fig. S7A,B). Our composition data during streptomycin treatment are qualitatively consistent with some previous studies (Stecher et al., 2007), but differ from others (Thompson et al., 2015), perhaps reflecting administration- or microbiota-specific effects of antibiotics or housing conditions. These potential sources of variability underscore the value of controlled studies of antibiotic treatment in which only intended variables such as diet and housing vary. Our experimental design has allowed us to robustly identify the key role of diet (Fig. 5) and co-housing (Fig. 7) in the microbiota response to antibiotics.

Our findings have a wide range of translational implications. The error-correction methodology implemented in DADA2 for 16S analysis provides a stark view of the precarious loss in diversity experienced by certain taxa during antibiotic treatment, sometimes down to 2 or fewer ASVs such as in the case of *Bacteroides* or S24-7 (Fig. 2G-I), in contrast to OTU-picking based methods (Caporaso et al., 2010) that suggested that hundreds of species remained (Fig. S2G) and did not reveal the dominance of *B. vulgatus* during antibiotics that we observed both with DADA2 as well as the sequencing of isolates during antibiotic treatment (Fig. 5H). There is a seemingly permanent loss of diversity after antibiotics (Fig. 2, S3, 5), reflecting the extinction of sensitive Bacteroidetes species. Transplants of these species may hasten microbiota recovery after antibiotic treatment; for example, *Bacteroides* are amenable to transfer to *Bacteroides*-lacking hosts (Fig. 6). Moreover, the loss of taxa may open niches for opportunistic pathogens, highlighting the importance of autologous fecal transplants and/or targeted microbial therapies to restore colonization resistance. Strikingly, a switch to a diet lacking MACs clearly sensitizes the microbiota to antibiotic treatment (Fig. 5), and may increase the potential for pathogen invasion. In future studies, it will be intriguing to probe the extent to which specific dietary interventions (e.g., high fiber) can improve resilience and recovery, similar to the reversal of a persistent *Clostridium difficile* infection (Hryckowian et al., 2018). Interestingly, our experiments suggest that while antibiotics free up niches for colonization, they can also recalibrate microbiotas for future treatments, such as in H2 mice in which *Bacteroides* spp. increased after the first treatment, promoting rapid recovery during the second treatment (Fig. 6D,I).

To fully address the extent to which the microbiota can affect antibiotic response, many further studies will be required. Yet, our study highlights some striking similarities in the responses of conventional mice and humanized mice, such as the dominance during recovery of *B. vulgatus* strains in the two colonization states with identical V3V4 16S rRNA sequences. These data support the notion that human and mouse harbor enough overlap in microbiota composition and function that similarities in responses can be identified, as with osmotic perturbations (Tropini et al., 2018). Nonetheless, several lines of evidence suggest that conventional mice are more sensitive than humanized mice to ciprofloxacin treatment. First, there is essentially no recovery of CFUs/mL in conventional mice during treatment (Fig. 3E), potentially due to the lack of Bacteroidetes throughout treatment (Fig. 3F). Second, the decrease in alpha diversity was generally larger and more prolonged in conventional mice (Fig. S3D) than in humanized mice (Fig. 2B,S3B). Finally, the response to a second treatment mirrored the first treatment in conventional mice (Fig. 3E,F), whereas humanized mice displayed a more resilient response to a second treatment (Fig. 5B,C). While this resilience may be due to the emergence of antibiotic-resistant strains in the humanized microbiota after the first treatment, it is remarkable that family-level abundances remained virtually unchanged throughout the second treatment (Fig. 3F). The lack of complete immune function in humanized mice may partially account for the differences between humanized and conventional mice; it will be intriguing to investigate the extent to which Nonetheless, these findings suggest that human microbiotas may be more resilient to antibiotics than mouse microbiotas, consistent with the dramatic increase in disruption caused by ciprofloxacin in MD compared with STD conventional mice.

Antibiotic treatment has interesting commonalities with other common gut microbiota perturbations. Both antibiotics and osmotic diarrhea (Tropini et al., 2018) revealed sensitivity of the S24-7 family (Fig. 1G, 5D,G), its competition with Bacteroidaceae for niches, the key role of cage sanitation in recovery (Fig. 7), and transitioning to substantially different steady states (Fig. 3) (Tropini et al., 2018). There are also surprising similarities among the responses to streptomycin, ciprofloxacin, and rifaximin, despite the differences in their mechanisms of action and the resistance profiles of common gut commensals to these drugs (Fig. S4, S7A,B). Along with the common core at the point of minimal diversity across diets that support highly differing starting states (Fig. 5H), these findings suggest that the gut microbiota may have a stereotyped collapse mechanism, even when an insult decreases the bacterial load by many orders of magnitude (Fig. 1B). However, the increased heterogeneity among singly housed mice (Fig. 7) suggests that collapse and recovery may rely on a network of metabolic interactions that cannot be maintained when sufficient extinction has occurred. Our findings paint the picture of a microbial community that bends with global shifts in diet, antibiotics, and composition, yet are able to exhibit resilience if the community architecture can be reinforced through environment-mediated recolonization. Future studies investigating how microbiota respond to perturbations on multiple time scales should enable mapping of the landscape of community states and trajectories, empowering the design of resilient communities and reprogramming of dysbiosis.

## Acknowledgments

The authors thank Michelle St. Onge and Marina Grunina for helpful discussions and advice with media preparation, and Nassos Typas for careful readings of the manuscript. The authors acknowledge support from the Allen Discovery Center at Stanford on Systems Modeling of Infection (to K.M.N. and K.C.H.), the Stanford Center for Systems Biology under NIH grant P50-GM107615 (to K.C.H.). A.A.-D. is a Howard Hughes Medical Institute International Student Research Fellow and a Stanford Bio-X Bowes Fellow. M.F., J.L.S, and K.C.H. are Chan Zuckerberg Biohub Investigators. K.X. and R.A.O. acknowledge Fundação para a Ciência e Tecnologia for individual grants IF/00831/2015 and PD/BD/106000/2014.

## Methods

All mouse experiments were conducted in accordance with the Administrative Panel on Laboratory Animal Care, Stanford University’s IACUC. More information about antibiotic administration, dietary shifts, and sacrificing is provided in the SI. The SI also includes details regarding bacterial density and 16S quantifications, as well as strain isolation, antibiotic sensitivity measurements, and whole genome sequencing.

