## Supplemental methods and figures for "Recovery of the gut microbiota after antibiotics depends on host diet and environmental reservoirs"

***Mouse experiments***

All mouse experiments were conducted in accordance with the Administrative Panel on Laboratory Animal Care, Stanford University's IACUC. For humanized mouse experiments, mice were gavaged with a human fecal sample obtained from a healthy anonymous donor (American male living in the San Francisco Bay Area, CA, age 42, omnivorous diet) as previously described (Kashyap et al., 2013). The microbiota was allowed to equilibrate for 6-8 weeks before perturbation experiments commenced. For monocolonization experiments, mice were gavaged with an overnight culture of *Bacteroides thetaiotaomicron* and allowed to equilibrate for 2 weeks before streptomycin treatment commenced. Conventional Swiss-Webster mice (RFSW, Taconic) were bred from an in-house colony. Mice were orally gavaged with antibiotics (Sigma Aldrich) dissolved in 200 µL water for five days. For experiments involving a dietary switch, mice were first fed a standard diet (Purina LabDiet 5010) rich in MACs, and then a defined low-MAC diet (Harlan TD.86489) in which the sole carbohydrates are sucrose (31% w/w), cornstarch (31% w/w), and cellulose (5% w/w).

Mice were euthanized with CO_2_ and death was confirmed via cervical dislocation. After sacrifice, sections of the ileum, proximal colon, and distal colon were preserved, embedded, sectioned, stained and imaged as previously described (Earle et al., 2015).

***Quantification of bacterial densities***

Anaerobic and aerobic bacterial densities were quantified by spot plating on duplicate Brain Heart Infusion agar plates supplemented with 10% defibrinated horse blood. Aerobic plates were incubated in a 37 °C warm room and anaerobic plates were placed in a 37 °C incubator inside an anaerobic chamber (Coy Laboratory Products).

***16S rRNA analyses***

DNA was extracted from whole fecal pellets with the PowerSoil and PowerSoil-htp kits (MO BIO, San Diego, CA). 16S rRNA amplicons were generated using the Earth Microbiome Project-recommended 515F/806R primer pairs. PCR products were cleaned, quantified, and pooled using the UltraClean 96 PCR Cleanup kit (MO BIO) and Quant-It dsDNA High Sensitivity Assay kit (Invitrogen). Samples were sequenced with 250- or 300-bp reads on a MiSeq (Illumina).

Samples were de-multiplexed and analyzed using DADA2 and QIIME v. 1.8 as previously described (Callahan et al., 2016; Caporaso et al., 2010). Custom MATLAB (MathWorks) scripts were used to analyze OTU distributions during antibiotic timecourses. Glycoside hydrolase (GH) imputations were performed as previously described (Zhang et al., 2018)), on 11/19/2018 using the dbCAN2 meta server. Collections of CAZy types were visualized using Anvi’o v. 5.2 (Eren et al., 2015). Anv'io was also used in conjunction with DIAMOND, MCL, and PyANI to perform gene clustering and calculation and visualization of ANI (Buchfink et al., 2015; Pritchard et al., 2016; van Dongen and Abreu-Goodger, 2012).

***Isolation of strains***

For isolation of *Bacteroides* strains, frozen mouse fecal samples were resuspended in PBS and plated onto Bacteroides Bile Esculin agar/ Brucella Laked Blood agar with kanamycin and vancomycin (BBE/LKV) plates (Anaerobe Systems). This medium selects for *Bacteroides* spp. Plates were incubated at 37 °C in an anaerobic chamber. After 2 days, colonies were grown in PYG (Anaerobe Systems) for 2 days and glycerol stocked.

**In vitro *measurements of antibiotic sensitivity***

All strains were grown anaerobically in a plate reader with constant shaking in pre-reduced BHIS (BHI broth supplemented with 0.5 µg/mL porcine hemin and 0.5 µg/mL vitamin K1). After 24 h of growth, cultures were diluted 100-fold in fresh medium and grown for another 24 h. Cultures were then diluted 200-fold into fresh medium containing antibiotics (ciprofloxacin, streptomycin, and rifaximin) at various concentrations in 384-well plates and grown at 37 °C. After 48 h of growth, absorbance was measured in a plate reader. The minimum dose of antibiotic at which absorbance was indistinguishable from media controls was used to determine the MIC.

***Preparation of mouse fecal pellets for mass spectrometry analysis***

Fecal samples were taken from mice at indicated times and stored at -80 °C until the time of mass spectrometry analysis. Fecal pellets were placed in tared, 2 mL mass spectrometer compatible tubes (USA Scientific, cat. # 1620-2700). 200 µL of methanol were added to each pellet. A set of 2.8 mm zirconium oxide ceramic beads (VWR, cat. # 10144-494) were added to each tube and samples were disrupted for 20 min at a frequency of 20 Hz on a TissueLyser II (Qiagen, cat. # 85300). After disruption, supernatants were obtained via centrifugation of samples for 15 min at 16,300*g*. 10 µL of cleared supernatant were removed and added to 10 µL of water + 0.1 % formic acid, mixed well, and placed in glass vials for analysis via mass spectrometry. If analysis was not immediately performed, isolated samples were stored at -20 °C.

***UPLC/electrospray ionization-MS analysis***

The LC-MS analysis was performed on an Agilent 6530 Accurate Mass QToF LC/MS with a dual electrospray ionization source (Agilent) connected to a Agilent 1290 Infinity II UPLC front-end equipped with an SB C18 column (1.8 µm, 3.0x100 mm, Agilent cat. #828975-302). The gas temperature was set to 350 °C. The VCap was set to 4000 V. The instrument was operated in positive-ion mode under MS conditions. The absorbance threshold was set at 5000 and the relative threshold was set at 1%. Data were collected and stored in centroid form. The mobile phase system was made up of solvent A (water + 0.1% formic acid) and solvent B (95:5 acetonitrile:water + 0.1% formic acid), with a flow rate of 0.2 mL/min. The method and gradient used to elute the fecal extracts was: 0 min to 2 min 0% B, 2 min to 12 min 0% B to 100%, 12 min to 13 min held at 100% B, 13 min to 14 min 100% B to 0% B, 14 min to 15 min 0% B to re-equilibrate the column. All gradients were linear. The first 2 min of the run were sent to waste to remove salt. MS data were collected from 1.2 min until 12.5 min. 2 µL of each sample were injected/run. Analysis was performed using the Agilent MassHunter Qualitative Analysis Software. Exact masses were calculated using ChemDraw and analyzing standards. TICs were extracted using these exact masses ±100 ppm, the preset default by MassHunter to generate EICs. EICs were manually integrated and the area under the peak was calculated by MassHunter. Area under the curve was then adjusted to be per gram of fecal matter input during fecal preparations.

**Supplemental Figure Legends**

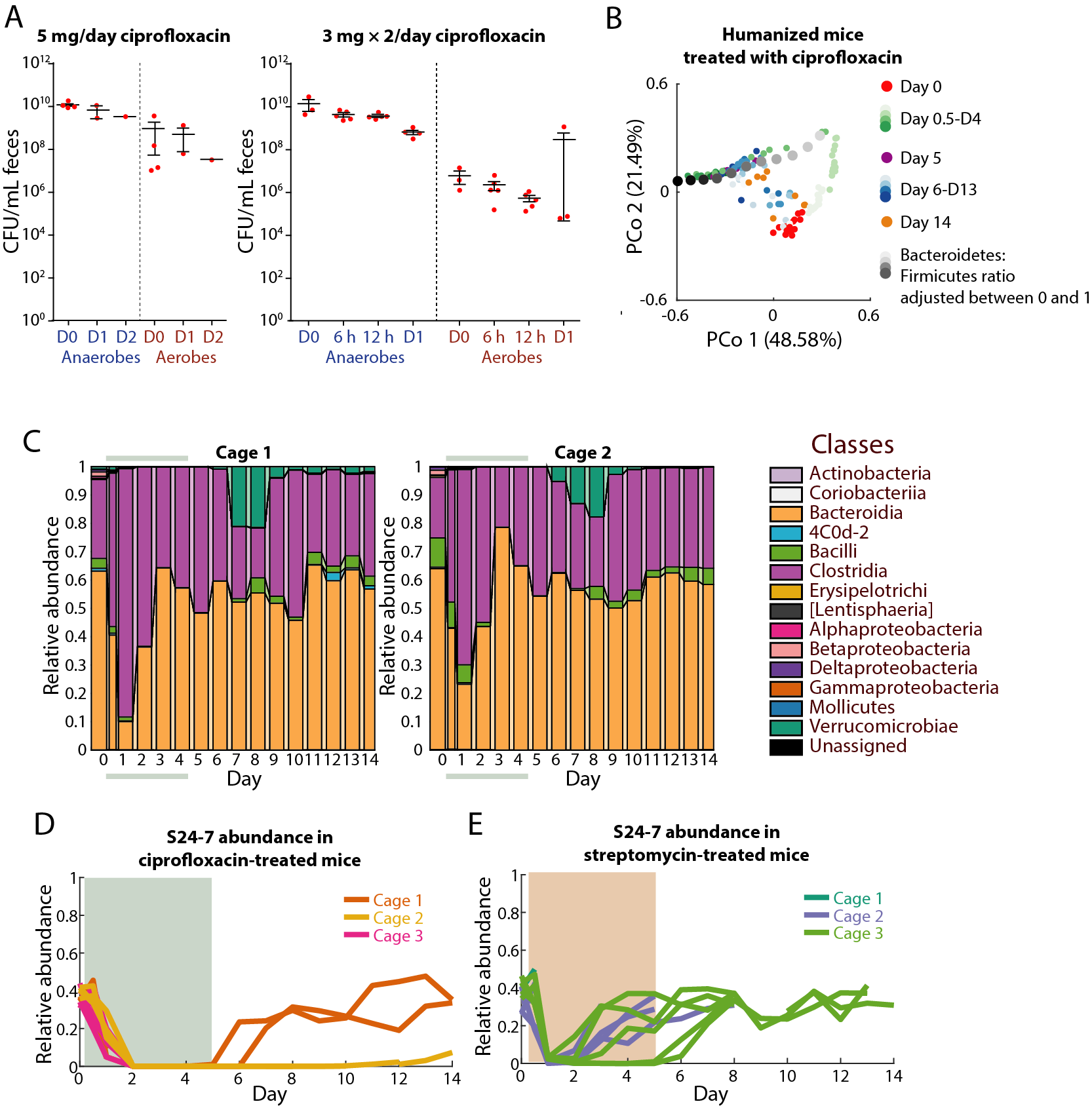

**Figure S1: Analyses of streptomycin- and ciprofloxacin-treated mice.**

A) Culturable bacterial densities in feces of mice treated with 5 mg ciprofloxacin/day or 3 mg twice daily revealed a larger effect with twice daily gavaging.
B) Computational adjustment of the Bacteroidetes:Firmicutes ratio through global rescaling of the fractional abundances of each phylum resulted in movement along principal coordinate 1.
C) Class-level community composition in feces in two cages of ciprofloxacin-treated humanized mice demonstrates insufficiency of class-level analyses to detect cage-specific differences in composition.

D,E) Relative abundance of S24-7 during (D) ciprofloxacin and (E) streptomycin treatment of humanized mice.

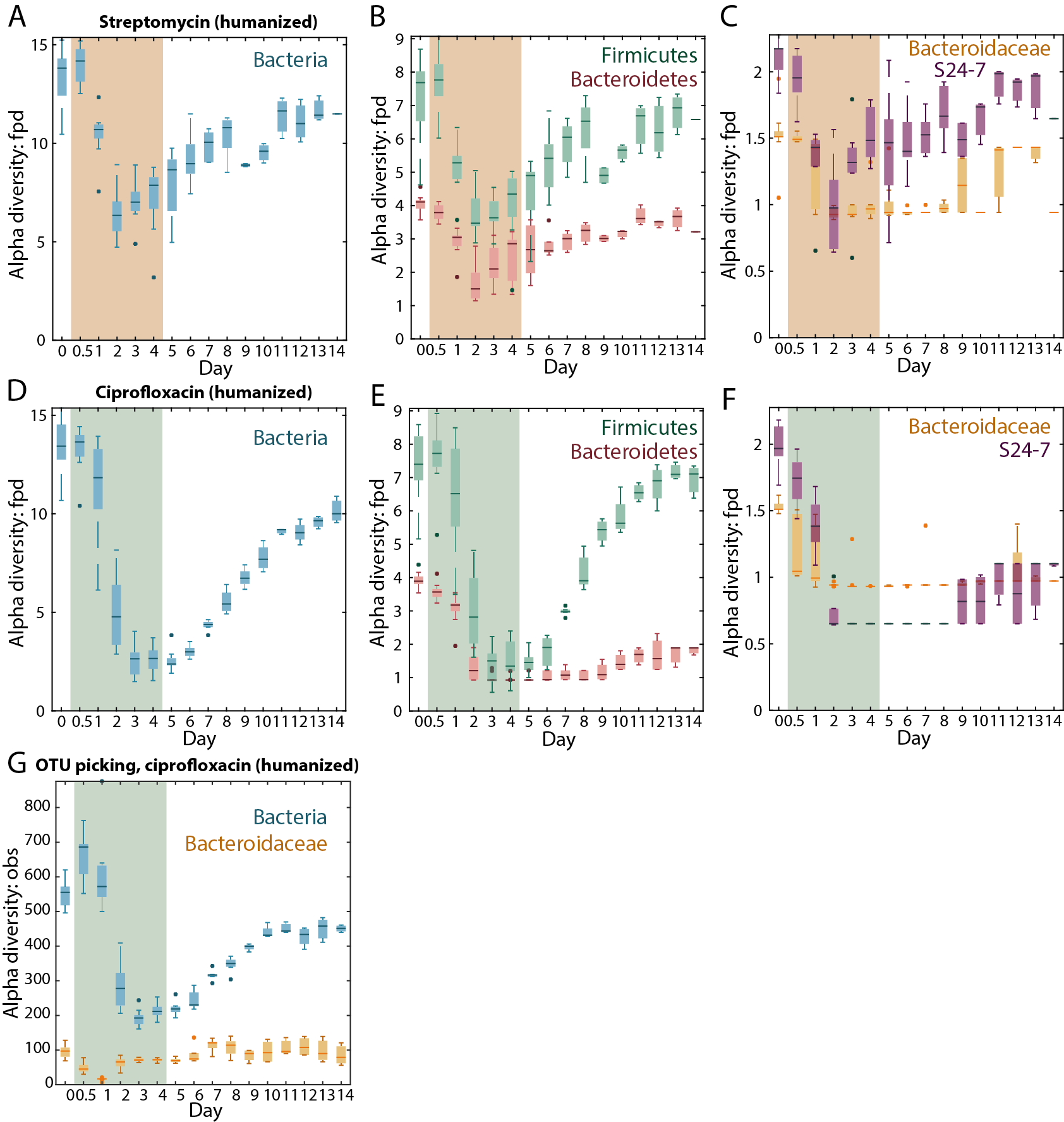

**Figure S2: Diversity analyses at different taxonomic levels for streptomycin- and ciprofloxacin-treated humanized mice.**

A,D) Alpha diversity (Faith's PD) of all bacteria in (A) streptomycin-treated humanized mice and (D) ciprofloxacin-treated humanized mice. Antibiotic treatment period marked in colored area.

B,E) Alpha diversity (Faith's PD) of phyla Firmicutes (green) and Bacteroidetes (pink) in (B) streptomycin-treated humanized mice and (E) ciprofloxacin-treated humanized mice. Antibiotic treatment period marked in colored area.

C,F) Alpha diversity (Faith's PD) of families S24-7 (purple) and Bacteroidaceae (yellow) in (C) streptomycin-treated humanized mice and (F) ciprofloxacin-treated humanized mice. Antibiotic treatment period marked in colored area.

G) Alpha diversity (observed ASVs) of ciprofloxacin-treated humanized mice calculated using *de novo* OTU-picking instead of DADA2.

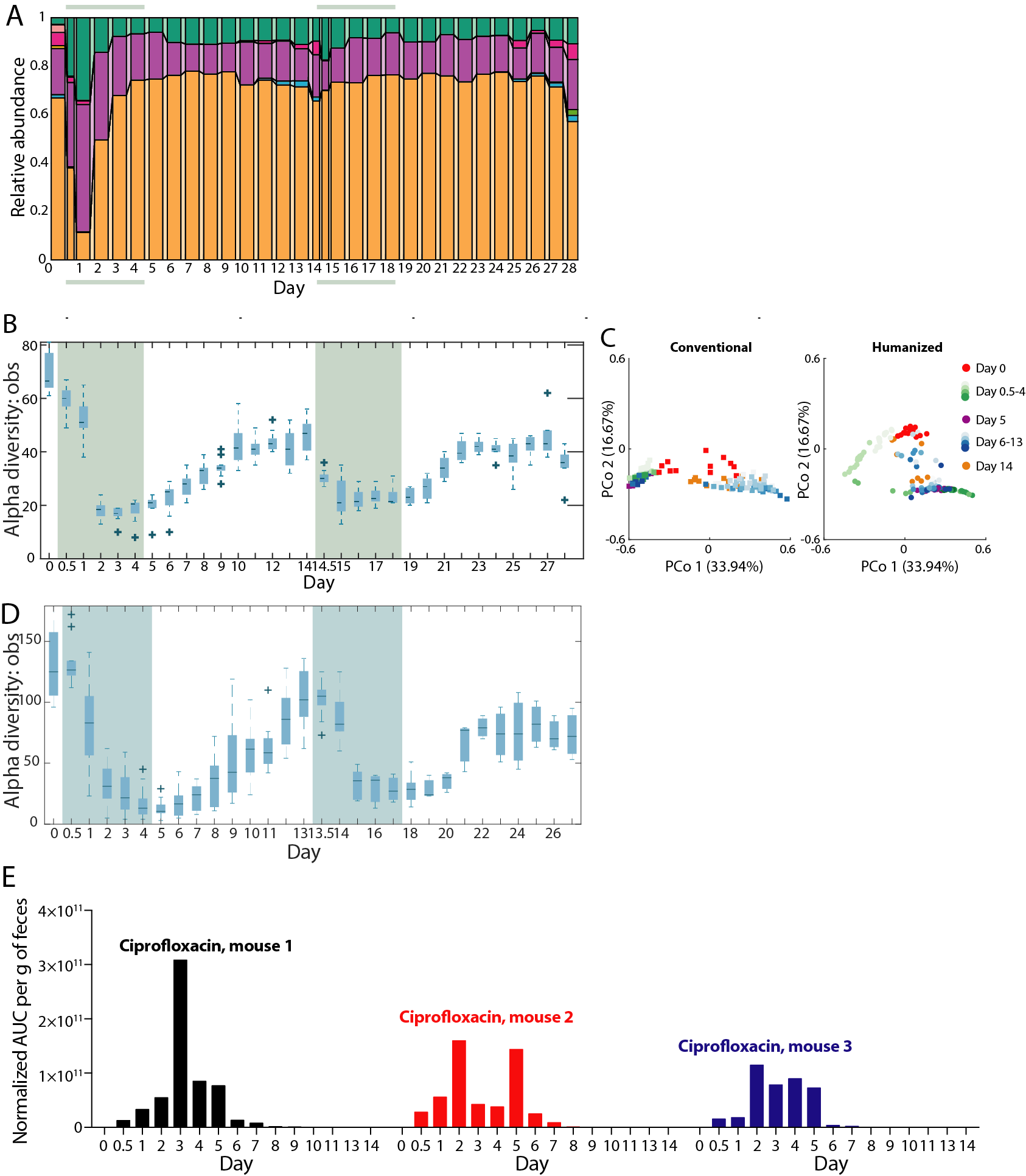

**Figure S3: Comparisons of humanized and conventional mice during double treatment with ciprofloxacin, and mass spectrometry analyses of ciprofloxacin-treated mice.**

A) Class-level community composition in feces of humanized mice further demonstrated the robustness of humanized mice to the second treatment.

B,D) Alpha diversity (observed ASVs) of all bacteria in (B) humanized and (D) conventional mice. The diversity decreased less in the humanized population during the second treatment than during the first.

C) Principal coordinates analyses (PCoA) of community composition in conventional and humanized mice during ciprofloxacin treatment reveal a conserved trajectory. Analyses used weighted UniFrac distances (Methods).

E) Normalized concentrations of ciprofloxacin in feces of conventional mice treated with ciprofloxacin.

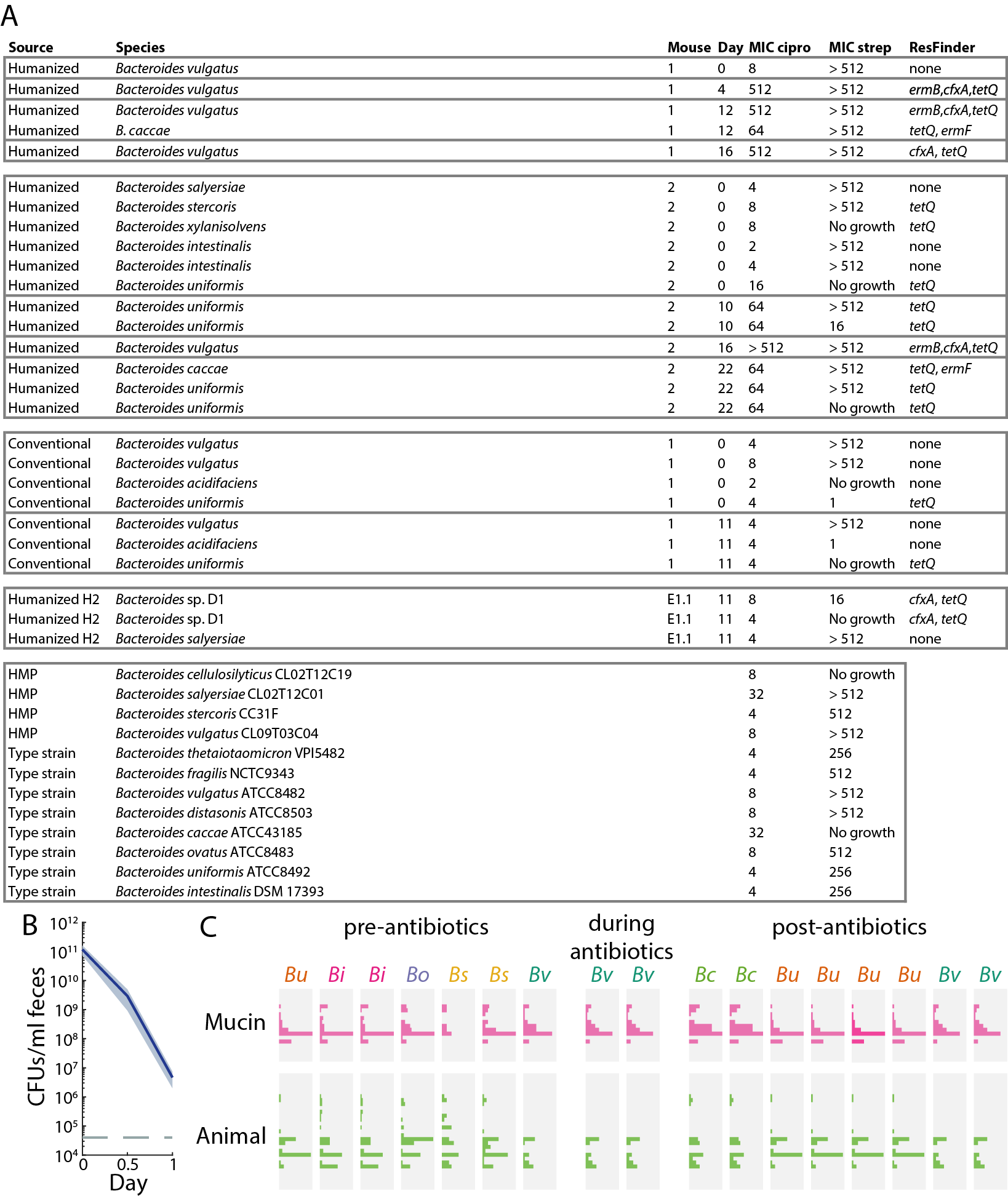

**Figure S4: MIC measurements and glycoside hydrolase of *Bacteroides* spp.**

A) Table of ciprofloxacin and streptomycin MICs for representative *Bacteroides* isolates and type strains, as well as any antibiotic resistance genes detected using ResFinder (<https://cge.cbs.dtu.dk/services/ResFinder/>).
B) Culturable anaerobic fecal densities in ex-germ-free mice (*n* = 10) colonized with *B. thetaiotaomicron* for 2 weeks prior to treatment with 20 mg streptomycin on day 0 and day 1. Bacterial loads collapsed as in humanized mice (Fig. 1B). Error bars: standard error of the mean.
C) Presence of carbohydrate active enzymes in *Bacteroides* spp*.* isolated from humanized mice. Mucin enzymes were over-represented in *Bc* post-antibiotics.

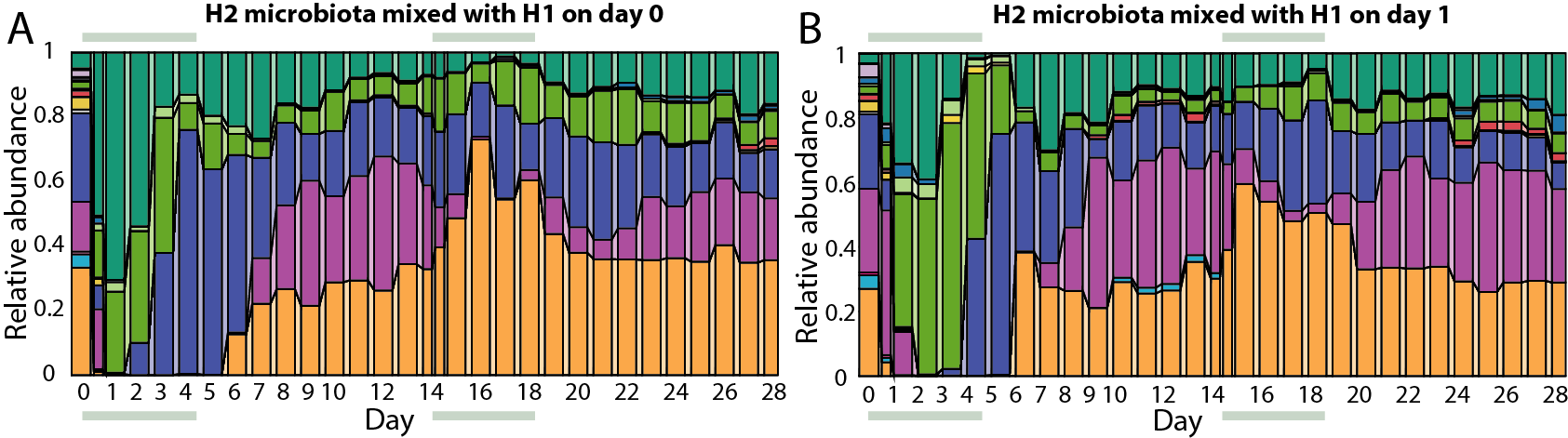

**Figure S5: Analyses of microbiota dynamics during cross-housing experiment.**

A,B) Family-level community composition in feces of H2 mice mixed with H1 mice on (A) day 0 and (B) day 1.

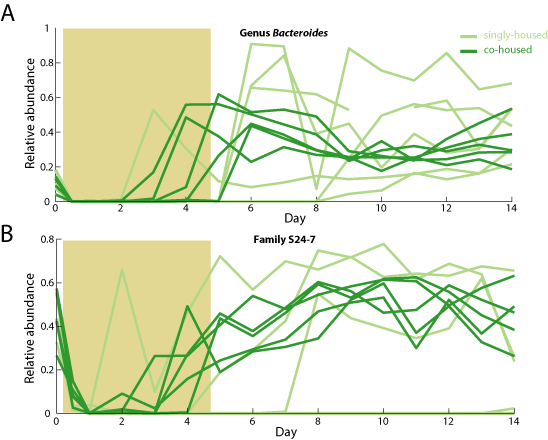

**Figure S6:** **Trajectories of Bacteroidetes taxonomic classes in singly housed and co-housed conventionalized mice.**

A,B) Relative abundance of the (A) *Bacteroides* genus and (B) S24-7 family in singly- and co-housed conventional mice. Antibiotic treatment period is denoted by colored rectangle. Recovery of both families in more heterogeneous in singly housed mice.

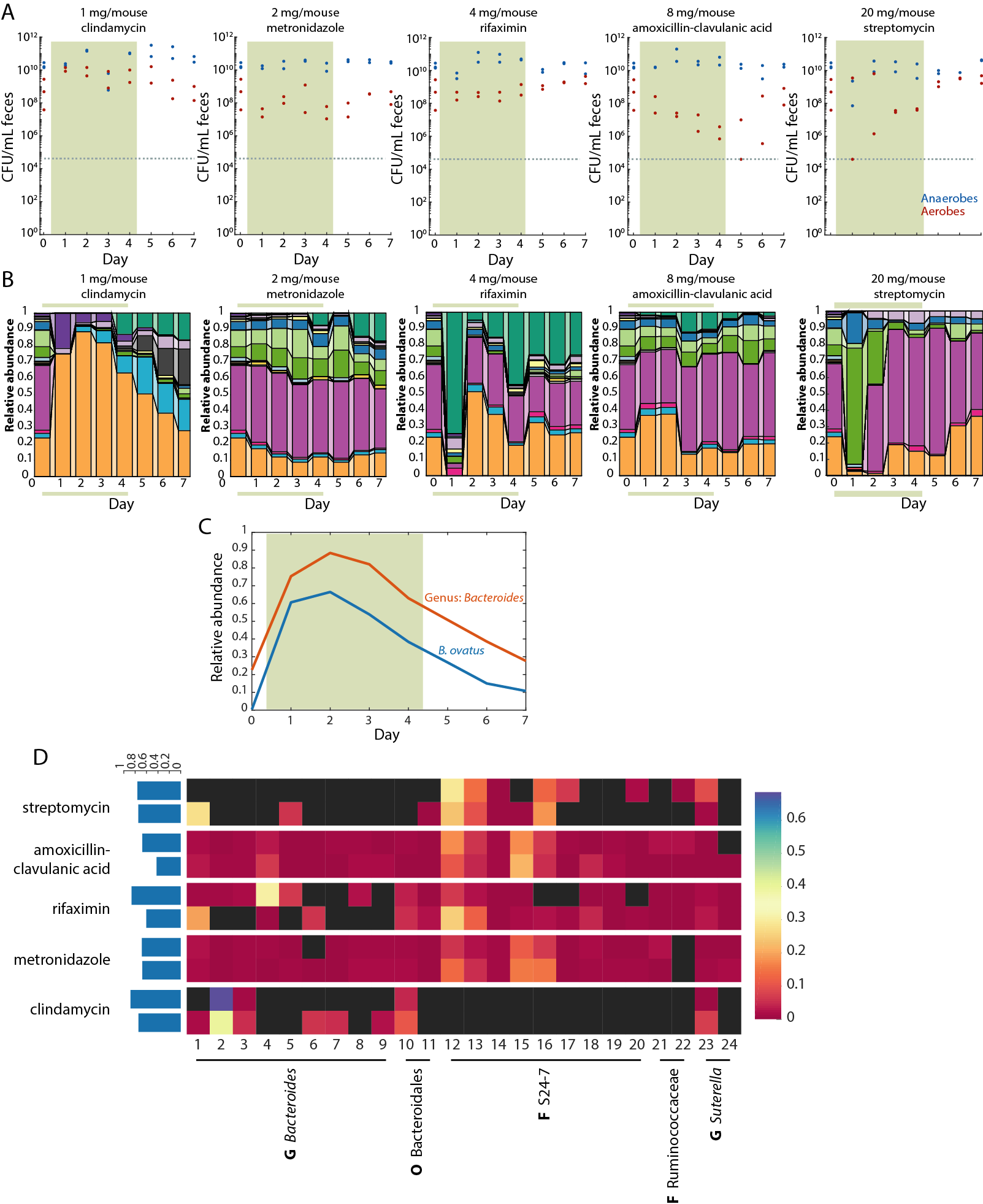

**Figure S7: Response of humanized mice to a wide range of antibiotics.**

A) Culturable anaerobic (blue) and aerobic (red) fecal densities in humanized mice gavaged with antibiotics for 5 days.
B) Family-level community composition in feces of mice in (A).
C) Relative abundance of a *B. ovatus* ASV and the genus *Bacteroides* in clindamycin-treated mice reveals domination of *B. ovatus*.
D) A conserved core microbiota of 24 species is observed in the 10 non-ciprofloxacin treated mice across antibiotics, largely composed of *Bacteroides* and S24-7 species.
